# Seagrass Extent Expansion in the Gulf of Mexico and Northwestern Caribbean (1987–2021)

**DOI:** 10.1101/2025.11.10.687714

**Authors:** Luis Lizcano-Sandoval, Susan Bell, Sergio Cerdeira-Estrada, Beatriz Martinez-Daranas, Enrique Montes, Brigitta van Tussenbroek, Frank Muller-Karger

## Abstract

Historically, the areal extent of seagrass beds in the Gulf of Mexico, Florida peninsula, and Northwestern Caribbean (Mexico and Cuba) have been estimated to cover approximately 35,510 km^2^. However, the absence of regular monitoring limits our ability to detect interannual variability and up-to-date health conditions, and to respond to stressors, such as degraded water quality. Seagrass areal extent is one of the core Essential Ocean variables (EOV) of the Global Ocean Observing System (GOOS). Resource managers and researchers are interested in how this seagrass extent and the distribution of this habitat changes over time to focus possible action to mitigate impacts of various stressors, such as degraded water quality. In this study, we mapped 19,405 km^2^ of seagrasses for the region for the period 2019–2021, using satellite imagery from Landsat 5, Landsat 7, Landsat 8, and Sentinel-2. Interannual changes in seagrass areal extent at 17 representative sites indicated that seagrass habitats increased by 12.3% between 1987 and 2021. Sites off the West Florida coast showed the highest increasing rates in seagrass extent (up to 3.1% yr^-1^) and lowest rate (-0.43% yr^-1^) was observed in Biscayne Bay, South Florida. Seagrasses in the Mexican Caribbean showed positive extent expansion rates (1.3–1.8% yr^-1^), while in Texas were stable over the period of study. Continuous observations of seagrasses like these help managers to understand spatial and temporal patterns of change and encourage further studies on seagrass ecology and carbon stocks.

## 1. Introduction

Seagrasses are marine angiosperms that are distributed widely in shallow coastal systems of the world (Short et al., 2007). Seagrasses in coastal waters of the Gulf of Mexico (also named Gulf from hereon) are often exposed to conditions that may be unfavorable for photosynthesis and growth, including turbidity and pollution (Yarbro and Carlson, 2016), drought and extreme salinity events (up to 50–70 salinity units) (Handley et al., 2007; Wilson and Dunton, 2018), and high sedimentation (Carter et al., 2011; Handley et al., 2007; Pham et al., 2014). Many seagrass beds in the Caribbean Sea, such as along the Mexican coast, develop under typically nutrient-limited conditions. However, nutrient input events may take place from submarine underground water discharges and sargassum beaching events (Gómez et al., 2022; van Tussenbroek, 2011). Tropical storms and extreme high and low temperature periods also have direct repeated and cumulative impacts on seagrasses of the region (Carlson et al., 2010; Kowalski et al., 2018; Lucas and Carter, 2013; Tomasko et al., 2020; van Tussenbroek et al., 2014).

Seagrass areal extent is one of the core Essential Ocean variables (EOV) of the Global Ocean Observing System (GOOS) (Duffy et al., In review). The areal extent of global seagrass habitats is highly uncertain, and is estimated to range from 177,000 to 600,000 km^2^ (Duarte et al., 2010; McKenzie et al., 2020). Seagrasses have long been thought as declining globally (Waycott et al., 2009), but it is difficult to make such assessments when the uncertainty in areal extent of these habitats is so large. More locally, long-term recovery has been observed in locations around Europe (de los Santos et al., 2019) and in estuaries such as Tampa Bay in the United States (Lizcano-Sandoval et al., 2022; Sherwood et al., 2017), generally due to effective management and policy efforts informed by monitoring data. However, monitoring programs over regional and larger scales are limited (Handley et al., 2007; van Tussenbroek et al., 2014; Yarbro and Carlson, 2016).

In coastal areas of the Gulf of Mexico, the Mexican Caribbean, and Cuba seagrass meadows have been estimated to extend over an area of 35,510 km^2^ (UNEP-WCMC and Short, 2021). Of this, approximately 40% is located around Cuba, 30% off the Florida peninsula, 20% in the Mexican Gulf states, 7% in the Mexican Caribbean, and 3% in the other US Gulf states. These values are mainly derived from comprehensive aerial surveys and visual interpretations supported by field observations and areas of occurrence, that at present has large geographic gaps in coverage and that does not allow assessments of variation over time (Handley et al., 2007; Handley and Lockwood, 2020; Yarbro and Carlson, 2016). Here we sought to address these limitations by (1) filling geographic gaps in seagrass distribution extent maps in the Gulf of Mexico, Florida peninsula, Mexican Caribbean, and around Cuba, (2) developing time series of annual seagrass extent in selected sites in the region and estimate rates of change, and (3) characterizing some of the potential drivers of seagrass change. To do this, we used extensive inventories of publicly-available satellite remote sensing data now available from the historical series of Landsat and Sentinel-2 satellites (Lizcano-Sandoval et al., 2025, 2022).

## 2. Methods

### 2.1. Study area

In order to develop our study of seagrass extent, the general location of these habitats was identified from the global seagrass distribution dataset of UNEP-WCMC and Short (2021), including descriptions from the original work developed by Green and Short (2003), and compilations of reported seagrass distributions in the US Gulf of Mexico and Florida (Handley et al., 2007; Yarbro and Carlson, 2016), Mexico (Herrera-Silveira et al., 2019, 2018) and Cuba (Martínez-Daranas et al., 2009; Martínez-Daranas and Suárez, 2018) (Figure 1). A total of 93 sites were inspected for potential seagrass mapping, but only 78 sites were selected for a regional comprehensive study of seagrass extent (Supplementary Table S1). Of these, 17 sites (15 sites in USA and 2 in Mexico) were selected for annual time series assessment (1987–2021) (Figure 1). The complexity for seagrass mapping in Cuba due to benthic habitat heterogeneity, water quality (i.e., turbidity), and scarce historical seagrass data did not allow time series mapping. Satellite imagery was visually inspected to select sites based on the following criteria: 1) seagrass meadow presence has been reported in the literature; 2) seagrass meadows are assessable under Sentinel-2 imagery (10 m per pixel) or Landsat imagery (30 m per pixel); 3) the site is located in optically shallow waters; and 4) optical water quality (i.e., transparency) allows the observation of the bottom from satellite imagery at least once and ideally a few times per year. Sites that did not meet these criteria were discarded. Most of the seagrasses in this region grow in shallow areas (<5 m depth), with few exceptions in the Mexican Caribbean and Yucatan Peninsula reported up to ∼10 m depth (de Almeida et al., 2022; Palafox-Juárez and Liceaga-Correa, 2017; Pérez Espinosa et al., 2019), Cuba up to ∼24 m depth (Martínez-Daranas and Suárez, 2018), and the Southwest Florida shelf up to 30 m depth (Handley et al., 2007).

**Figure 1.**
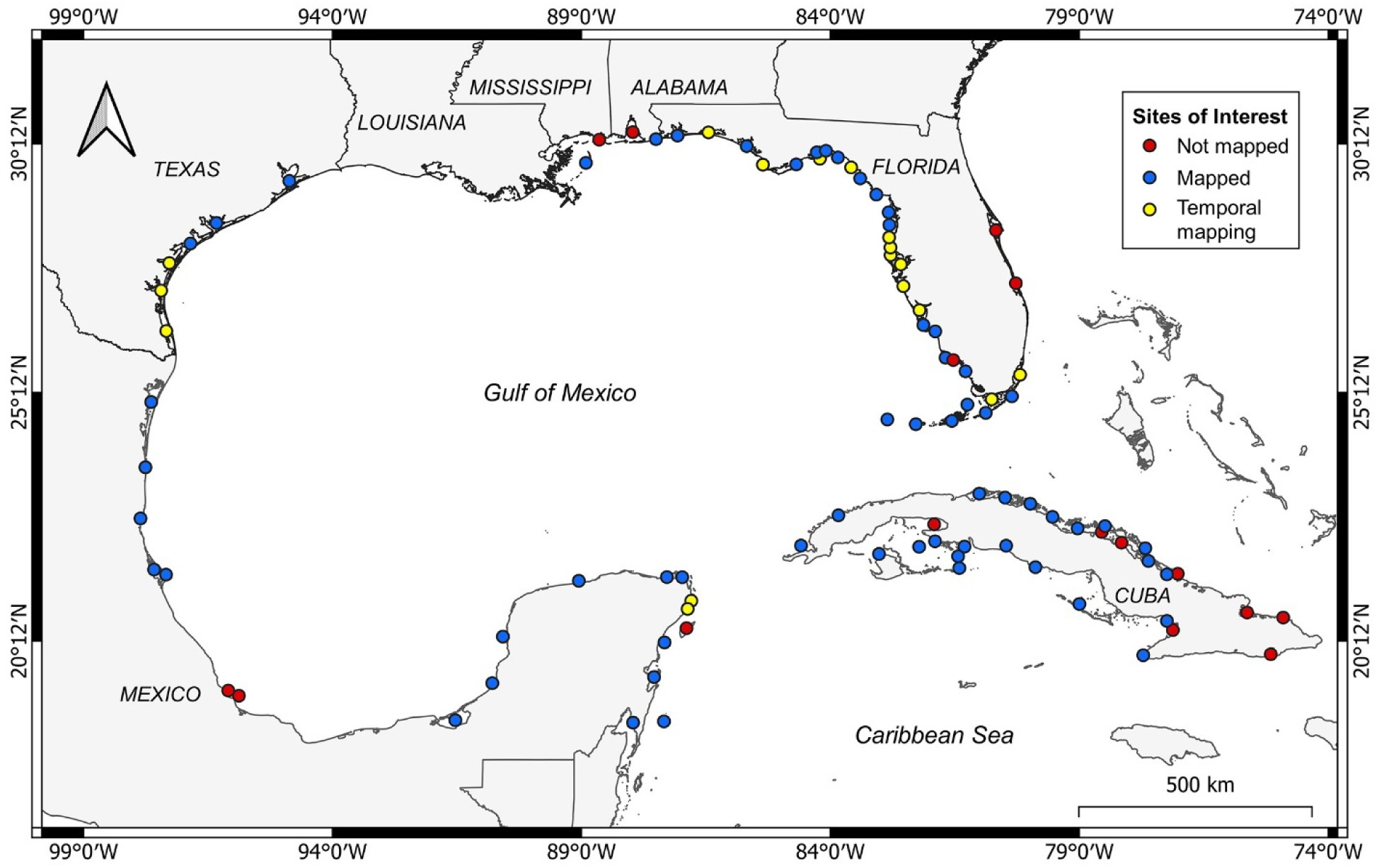
Sites of interest for satellite seagrass mapping identified from seagrass presence reports. Sites selected for seagrass extent mapping (2019–2021) are shown both in blue and yellow. The yellow dots represent those sites selected for time series mapping (1987–2021). The sites with reported seagrass that were not possible to map due to turbidity and sparse seagrass beds are indicated by red dots.

### 2.2. Satellite seagrass mapping

To develop a general baseline map of seagrasses of the region of study, we combined extent maps derived from Sentinel-2 (A and B) Multispectral Instrument (MSI) imagery corresponding to years for which we collected the most comprehensive ground truth data and processed it. This was from 2019 for Florida; 2020 for Louisiana, Texas and Mexico; and 2021 for Cuba (Supplementary Table S1). To generate the multi-decadal time series for the 17 sites, imagery from several satellite sensors was used. This included, from most recent to oldest: Sentinel-2 for 2016–2021; Landsat 8 Operational Land Imager (OLI) for 2013–2016; Landsat 7 Enhanced Thematic Mapper (ETM+) for 1999–2002; and Landsat 5 Thematic Mapper (TM) for 1987–2012. Landsat 5 was launched in 1984, but usable and more available images for the region were only found starting in 1987, for US territory. For other sites in the Mexican coast, Landsat 5 images were only available between 1993 and 2002. The data gaps in Landsat 5 outside US territory is due to non-copied data to the United States Geological Survey (USGS) archive (Kovalskyy and Roy, 2013). For instances when Sentinel-2 images from 2016 were not available on any of the sites, Landsat 8 images were used instead. Sentinel-2A was launched in June 2015, and Sentinel-2B was launched in March 2017. Therefore, more images became available after 2017. Sentinel-2 imagery was preferred over Landsat 8 due to the higher spatial resolution of the Sentinel-2 sensors (10 m pixels compared to the 30 m for Landsat (Supplementary Table S2). All imagery was accessed and processed through Google Earth Engine (GEE), a cloud-based platform that offers access to multi-petabyte satellite imagery and to high-performance computing resources (Gorelick et al., 2017).

The images for the study were initially selected using a series of criteria. We used the mean value of the Normalized Difference Turbidity Index (NDTI) to evaluate whether water quality permitted us to derive any meaningful seagrass maps. The NDTI uses the near-infrared (NIR) and short-wave infrared (SWIR) bands to evaluate water quality for a specific area of the image. Cloud cover percentage and the number of valid pixels remaining in the imaged seagrass region, after land and cloud masking (Lizcano-Sandoval et al., 2025, 2022). This process was repeated for each site of interest (i.e., a cluster of pixels in a polygon defined loosely around the regions where there would be seagrasses). If two or more sites were located inside one satellite image, each site was evaluated for adequate atmospheric and water quality conditions. After selecting satellite images, we followed the workflow for seagrass mapping described by Lizcano-Sandoval et al. (2022) and Lizcano-Sandoval et al. (2025) (Supplementary Figure S1).

The study sites are usually optically shallow (<10 m depth), but any effect of depth on remote sensing reflectance was compensated by using the Depth-Invariant Index (DII) (Supplementary Figure S1) The DII uses the blue and green band ratio based on the reflectance of a particular bottom type, which depending on the site was sand or any other softbottom type, and the output layer is treated as an additional band of the respective satellite image (Green et al., 2000; Lyzenga, 1981). The DII was not applied in sites with frequent turbidity (e.g., Texas, Louisiana, Florida Bay) and a turbidity mask was implemented to improve image quality for mapping (Supplementary Figure S1). The turbidity mask was empirically developed by using adjustable threshold values of NDTI and the NIR band and applied for turbid sites (Supplementary Figure S2). The NIR to blue band ratio also helped to identify potential seagrass pixels and separate them from turbid pixels.

We collected extensive reference benthic habitat data working with collaborating groups, from the literature, or from open databases. More details about reference data are described in the section below. A Support Vector Machine classifier used 70% of the reference data points to train on the processed images. We used the coastal, blue, green, and red bands for Sentinel-2 and Landsat 8. For Landsat 5 and Landsat 7 we used the blue, green, and red bands (Supplementary Table S1). If a DII was run previously, then it was also included along with the optical bands as input for the classifier. The remaining 30% of the reference data was reserved for validation. The overall accuracy (OA), user’s and producer’s accuracies (UA, PA), and the Kappa statistic were estimated (Congalton and Green, 2009). The OA indicates how good the model is for classifying the different bottom types available. The PA and UA measure the performance of the model to classify each bottom type separately, from which we focused on the seagrass class. The validations were done for each classified image and for each annual seagrass mosaic.

The images classified for any one year were treated with a gaussian kernel of 3×3 pixels to reduce noise. The images were subsequently mosaicked together by keeping only seagrass pixels to produce what we define as an annual seagrass mosaic (Lizcano-Sandoval et al., 2022). Any patches that were visually identified as some other habitat, such as macroalgae, were masked out. This was necessary to have consistent estimates of seagrass extent and distribution for temporal comparisons; the results are therefore rather conservative.

The post-processed annual mosaics were uploaded to GEE as a final product that is open and free for public use (see Data Availability). Given the limitations of satellite remote sensing, image resolution, and any potential classification errors, the assumptions for the final product were: 1) it represent optically dense seagrass habitats in shallow waters (usually <10 m depth); 2) the sum of contiguous pixels represents an assessment of seagrass areal extent; 3) seasonal variations of seagrass shoot density were small and the stack of images provided the maximum annual extent observed; and 4) seagrasses were dominant in any seagrass-algae-coral mixed patches.

### 2.3. Georeferenced benthic habitat data

We used multiple historical field seagrass monitoring and distribution datasets to identify locations with seagrass meadows and collect reference data. First, we examined the world seagrass distribution dataset from the United Nations Environmental Programme World Conservation Monitoring Centre (UNEP-WCMC and Short, 2021). Occurrences of the most common seagrass species of the Northwestern Atlantic (*Thalassia testudinum*, *Halodule wrightii*, and *Syringodium filiforme*) between 1984–2021 from the Global Biodiversity Information Facility (GBIF) were also used as seagrass presence reference. We also examined data from regional and local monitoring programs, including the Fish and Wildlife Research Institute for Florida (FWRI), the Southwest Florida Water Management District (SWFWMD), and the Tampa Bay Estuary Program (TBEP) for West-Central Florida. Texas seagrass distribution data was obtained from the Galveston Bay Estuary Program, Texas Parks & Wildlife, and the Texas Seagrass Monitoring Program. The seagrass distribution datasets were complemented with habitat maps and descriptions from scientific literature (Supplementary Table S3). All these sources were used as reference to identify potential seagrass mapping sites in the region (Figure 1) and to collect respective georeferenced points.

Individual georeferenced points noting presence of a habitat type were preferred instead of polygons for training the classifier. Each point or pixel was treated as an independent observation assigned to a single class (seagrass, softbottom, or hardbottom), which was confirmed by visual interpretation of satellite images (Sentinel-2 and Landsat) to ensure correspondence before classification and increase accuracy. We used the Florida’s United Reef Map distributed by the FWRI (https://myfwc.com/research/gis/fisheries/unified-reef-map/), the Allen Coral Atlas maps (https://allencoralatlas.org/), and the Caribbean Benthic Habitat Maps 2021 distributed by The Nature Conservancy (https://sites.google.com/view/caribbean-marine-maps) to collect reference points of hardbottom. The softbottom points were mostly collected by visual interpretation. The baseline dataset for Florida was 2019; for Texas, Louisiana, and Mexico the ground truth was for 2020; and for Cuba it was 2021. These baseline datasets included a total of 33,909 points of seagrass presence distributed across the sites of study. Other points were assigned to softbottom (37,088) and hardbottom (3,547) classes. The softbottom class represented sand, mud, or any other bare softbottom type. The hardbottom class represented corals or rocks.

In order to derive multi-year time series at the 17 selected sites, we assumed that seagrass distribution was relatively stable over time in most areas; this facilitated reusing most of the corresponding baseline datasets per site to reconstruct reference datasets for other years and classify respective images. Points in our reference dataset were adjusted if changes in seagrass distribution were either indicated by the source data or visually detected in the corresponding satellite images for any one year. In total, approximately 608,000 georeferenced points were used for annual classifications and respective validations over 1987–2021 (50% seagrass, 47% softbottom, and 3% hardbottom).

### 2.4. Site characterization

A general characterization of 17 seagrass time-series sites was conducted using precipitation, sea surface temperature (SST), chlorophyll-a concentration, particulate organic carbon (POC), human population density, hurricane occurrences, and marine protected area (MPA) coverage, to find patterns that could relate to seagrass change rates. Precipitation data were sourced from the Climate Hazards Center InfraRed Precipitation with Station data (CHIRPS) at 5.5 km resolution and processed for inland areas corresponding to the nearest second-level administrative divisions (e.g. counties, municipalities, or equivalent) to each seagrass site. Annual precipitation data (mm yr^-1^) were obtained for 1987–2021, and mean annual precipitation (mm yr^-1^) data were computed for each site.

The SST (°C), POC (mg m^-3^), and chlorophyll-a (mg m^-3^) data were acquired from the Moderate Resolution Imaging Spectroradiometer (MODIS) Aqua satellite Level-3 products at 4.6 km resolution, provided by the NASA’s Ocean Biology Processing Group (https://oceancolor.gsfc.nasa.gov). These data were processed for 2003–2021, using a 10 km buffer around each seagrass site to incorporate surrounding pixels, and annual means were calculated for each variable. Additionally, the ETOPO1 bathymetry product at 1.8 km per pixel was used to mask shallow water pixels to reduce possible error estimations in ocean color measurements (e.g. chlorophyll-a, POC), due to bottom reflectance contamination. Pixels in less than 5 m depth were excluded from the calculations, in most of the sites. Florida Bay (East) was a very shallow and turbid site that was treated by removing pixels in less than 1 m depth, assuming that light in most of the site was poorly reflected off the bottom. The sites in the Mexican Caribbean are in clearer and deeper waters, which allowed to remove pixels in less than 10 m depth.

Population density data were derived from the Global Human Settlement Layer (GHSL) raster dataset (Pesaresi and Politis, 2023). This provided the number of inhabitants per 100 m grid point at 5-year intervals from 1975 onward. Data were processed for each seagrass site using a 10 km inland buffer to capture potential areas of influence on seagrass habitats. The overall mean of population density per site was obtained for 1985–2020.

Hurricane impacts were quantified using the Atlantic Hurricane Database (HURDAT2) of the National Hurricane Center (https://www.nhc.noaa.gov/data/#hurdat). Direct hurricane impacts were counted for 1987–2021 within a 20 km buffer around each seagrass site. Finally, Marine Protected Area (MPA) coverage at each seagrass site was estimated using the 2025 database from Protected Planet (https://www.protectedplanet.net).

A non-metric multidimensional scaling plot (NMDS) with Euclidean distance was used to visualize the ordination and similarity among seagrass sites based on the environmental and socio-economic variables. The significance of correlations between each variable and the two ordination axes was tested. This multivariate analysis was performed using the “vegan” package in R (version 4.5.1).

### 2.5. Trend analyses

Seagrass areal extent estimations were done for each area of study and used to evaluate trends at the 17 selected sites. The sites with historical seagrass extent data were grouped into five subregions: Northwest Florida (NWF: Big Bend A-BGBA, Big Bend E-BGBE, Choctawhatchee Bay-CHOC, St. Joseph Bay-STJB), West Florida (WF: Charlotte Harbor (North)-CHAN, Clearwater Harbor-CLEA, Sarasota Bay-SARA, Springs Coast-SPCC, St. Joseph Sound-STJS, Tampa Bay-TBAY), South Florida (SF: Biscayne Bay-BISC, Florida Bay (East)-FLBE), Texas (TX: Corpus Christi Bay-CCBA, Lower Laguna Madre-LOLM, Upper Laguna Madre-UPLM), and Mexican Caribbean (MXC: Benito Juarez-BJUA, Puerto Morelos-PMOR). Seagrass extent trends were based on linear regressions with non-zero intercept. Seagrass extent change per year (% yr^-1^) were estimated from slope values (km^2^ yr^-1^). The linear regressions met the assumptions of homoscedasticity. This approach was different from just evaluating trends based on trajectories estimated from initial and final values only. The multi-year data showed interannual variations and each annual point contributed to defining the linear trend. All plots were made in Python using the “matplotlib” package. Maps were made in QGIS (version 3.18.3).

## 3. Results

### 3.1. Regional seagrass distribution

In total, 623 satellite images spanning from 2019 to 2021 were classified across 78 sites to produce the regional seagrass distribution map of the Gulf of Mexico, Florida, Mexican Caribbean and Cuba (Figure 2). The total visible seagrass habitat extent across these regions was 19,405 km^2^. Of this total, Cuba represented 44.3%, the Florida peninsula 38.4%, Southern Gulf of Mexico (Mexico) 12.2%, the Northern Gulf of Mexico (Texas and Louisiana) 2.9%, and the Mexican Caribbean 2.1% (Supplementary Table S2). Regionally, 69% of the seagrass habitats in the USA were included in MPA’s. In Mexico and Cuba, MPA’s covered 85% and 42% of the seagrass habitats, respectively (Figure 2). The global median value for overall classification accuracy was 92%, with seagrass producer’s and user’s accuracies both at 95%, and a Kappa coefficient of 0.8 (Supplementary Figure S3).

**Figure 2.**
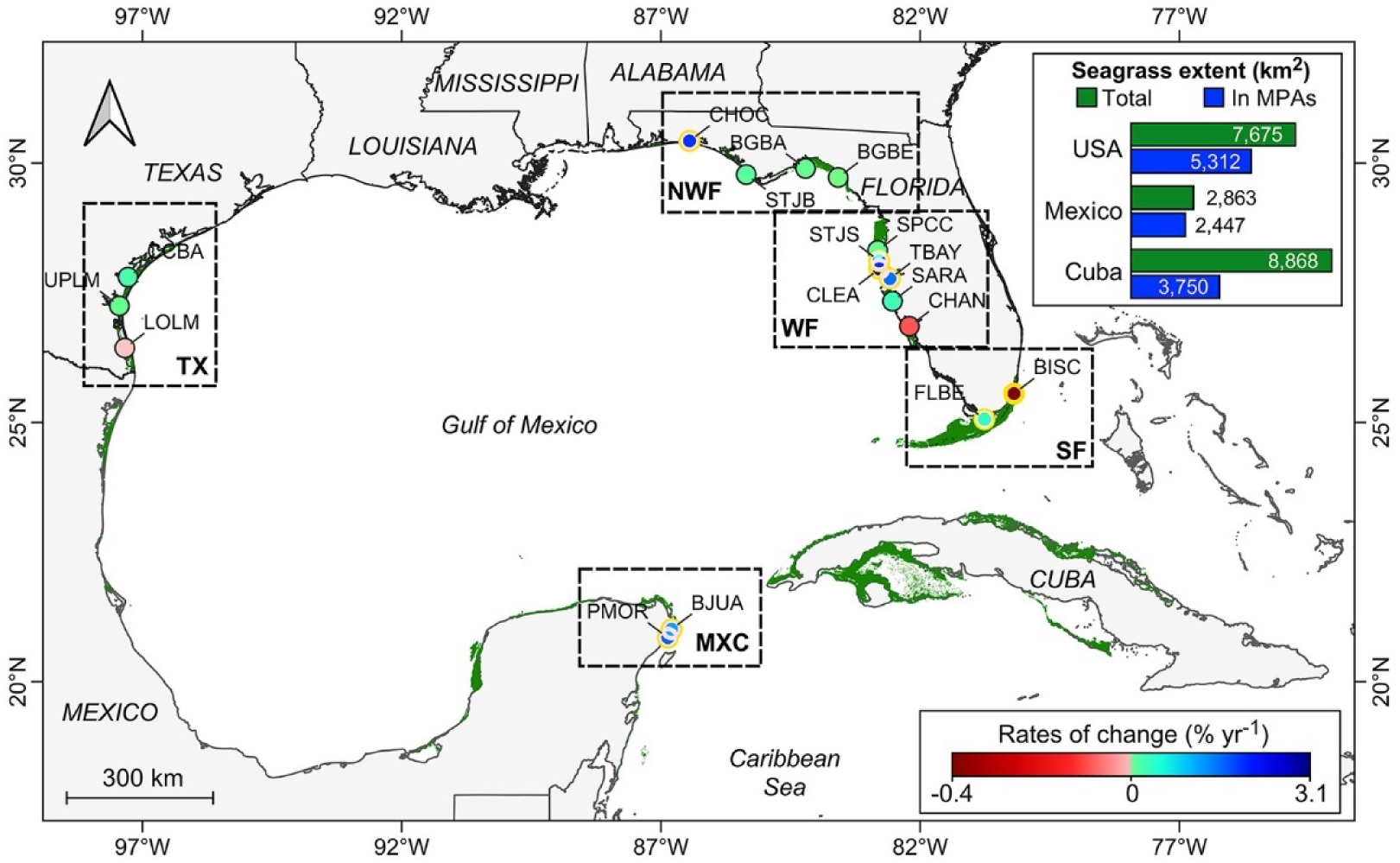
Regional seagrass distribution (in green) and rates of change (% yr^-1^) over 1987–2021 in 17 sites of South Florida (FL), West Florida (WF), Northwest Florida (NWF), Texas (TX), and Mexican Caribbean (MXC). The color of the circles indicates the seagrass extent rate of change per site according to the color bar. Sites with circles with yellow edges are statistically significant.

### 3.2. Long-term changes in seagrass areal extent

Time series of seagrass areal extent (Figure 3) were reconstructed from 1,811 classified images in the period 1987–2021 for 17 sites in five subregions: South Florida (SF), West Florida (WF), Northwest Florida (NWF), Texas (TX), and Mexican Caribbean (MXC). These sites represented 14% (2,633 km^2^) of the total seagrass mapped for the region in 2019–2021. Overall, the difference between the initial and final seagrass extent measurements across the 17 sites indicated an increase of 12.3% or 288 km^2^ in the period 1987–2021. Seagrass areal extent varied within this timeframe, but overall, seven sites showed significant increases in seagrass extent over time: Choctawhatchee Bay (2.04% yr^-1^), Clearwater Harbor (3.12% yr^-1^), St. Joseph Sound (2.05% yr^-1^), Tampa Bay (1.68% yr^-1^), Florida Bay (East) (0.58% yr^-1^), Benito Juarez (1.31% yr^-1^), and Puerto Morelos (1.79% yr^-1^). In contrast, Biscayne Bay showed significant decreases (-0.43% yr^-1^) (Figure 2, Figure 3, Table 1). Other no significant decreases in seagrass extent were observed for Charlotte Harbor (North) (-0.11% yr^-1^) and Lower Laguna Madre (-0.08% yr^-1^) (Table 1). The mean interannual variation in seagrass extent was usually not higher than 8.6%, except Choctawhatchee Bay, which showed a mean interannual variation of 20.9% (Table 1).

**Figure 3.**
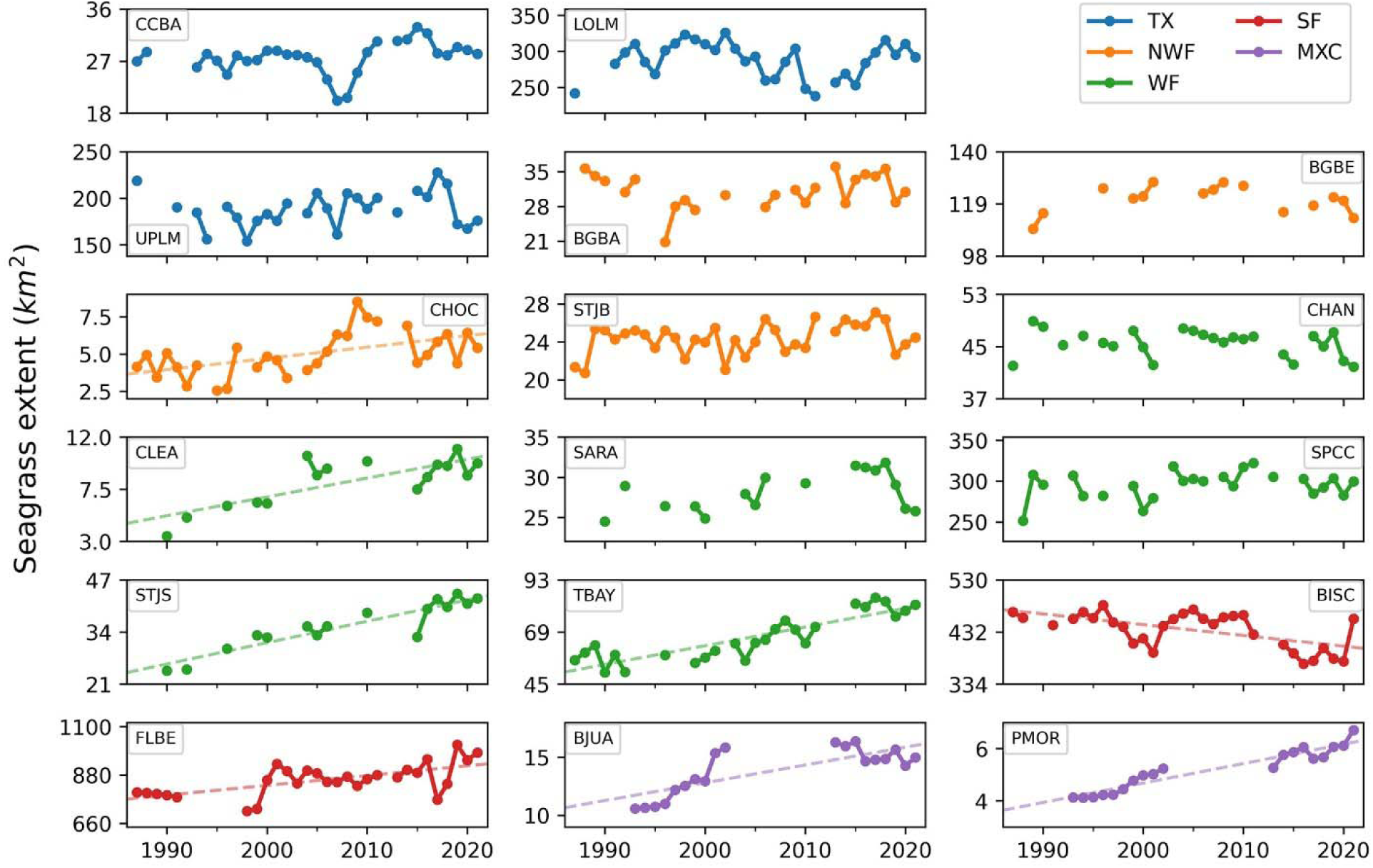
Seagrass extent at different sites of the Gulf of Mexico and Florida coasts. Texas (TX), Northwest Florida (NWF), West Florida (WF), South Florida (SF), and Mexican Caribbean (MXC). Trend lines are plotted for those sites with significant slopes (p < 0.05). See Table 1 for a description of site acronyms

**Table 1.**
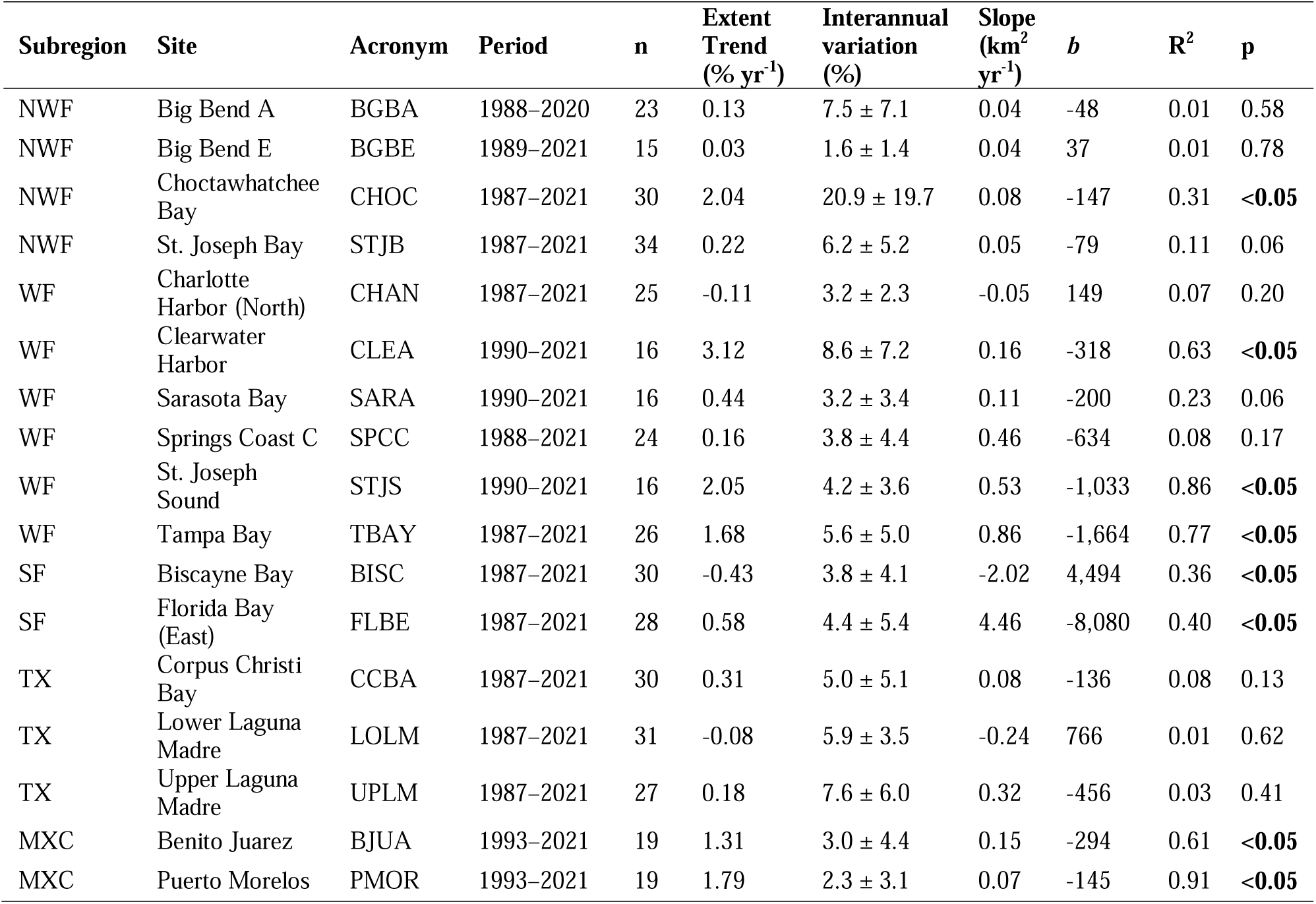
Temporal trends in visible seagrass areal extent in 17 sites of five subregions: Northwest Florida (NWF), West Florida (WF), South Florida (SF), Texas (TX), and Mexican Caribbean (MXC). The intercept (*b*), slope, R^2^, and p-value are the results of the linear fit. The mean *±* standard deviation of interannual variation (%) in seagrass extent over the respective period is also shown. The extent trends were derived from the slope values. Values of p <0.05 indicate significant linear regression.

Validation accuracies were usually higher than 70% and frequently observed around 90% in the overall accuracy (OA), user’s and producer’s accuracies (UA, PA) (Supplementary Figure S4). The Kappa statistic was usually higher than ∼0.5 and frequently observed above 0.8 (Supplementary Figure S4).

### 3.3. Site characterization

Mean values of the environmental and social characterization variables revealed gradients and patterns across seagrass sites (Figure 4). Ranges in these variables were as follows: population density of 0.1–15.3 inhabitants per 100 m pixel, precipitation of 631–1,629 mm yr^-1^, SST of 23.3–28.1°C, chlorophyll-a concentration of 0.1–12.2 mg m^-3^, POC of 43–869 mg m^-3^, hurricane frequency of 0–3 events, and MPA coverage of 0–100% (Supplementary Table S4). Subregional patterns were generally consistent among nearby seagrass sites, except for Florida Bay (East) and Biscayne Bay in South Florida, which exhibited distinct spatial ordination driven by contrasting population density, and chlorophyll-a and POC concentrations (Figure 4). All characterization variables showed significant correlations with the non-metric multidimensional scaling plot (NMDS) with Euclidean distance ordination axes (p < 0.05). The sites in Northwest Florida are mainly characterized by high concentrations of POC and chlorophyll-a and high precipitation. The Mexican Caribbean and Texas were more associated to high SST and hurricane impacts. High precipitation, high population density, and high MPA coverage were associated to some sites of the West and South Florida subregions in the vertical NMDS axis, (Figure 4).

**Figure 4.**
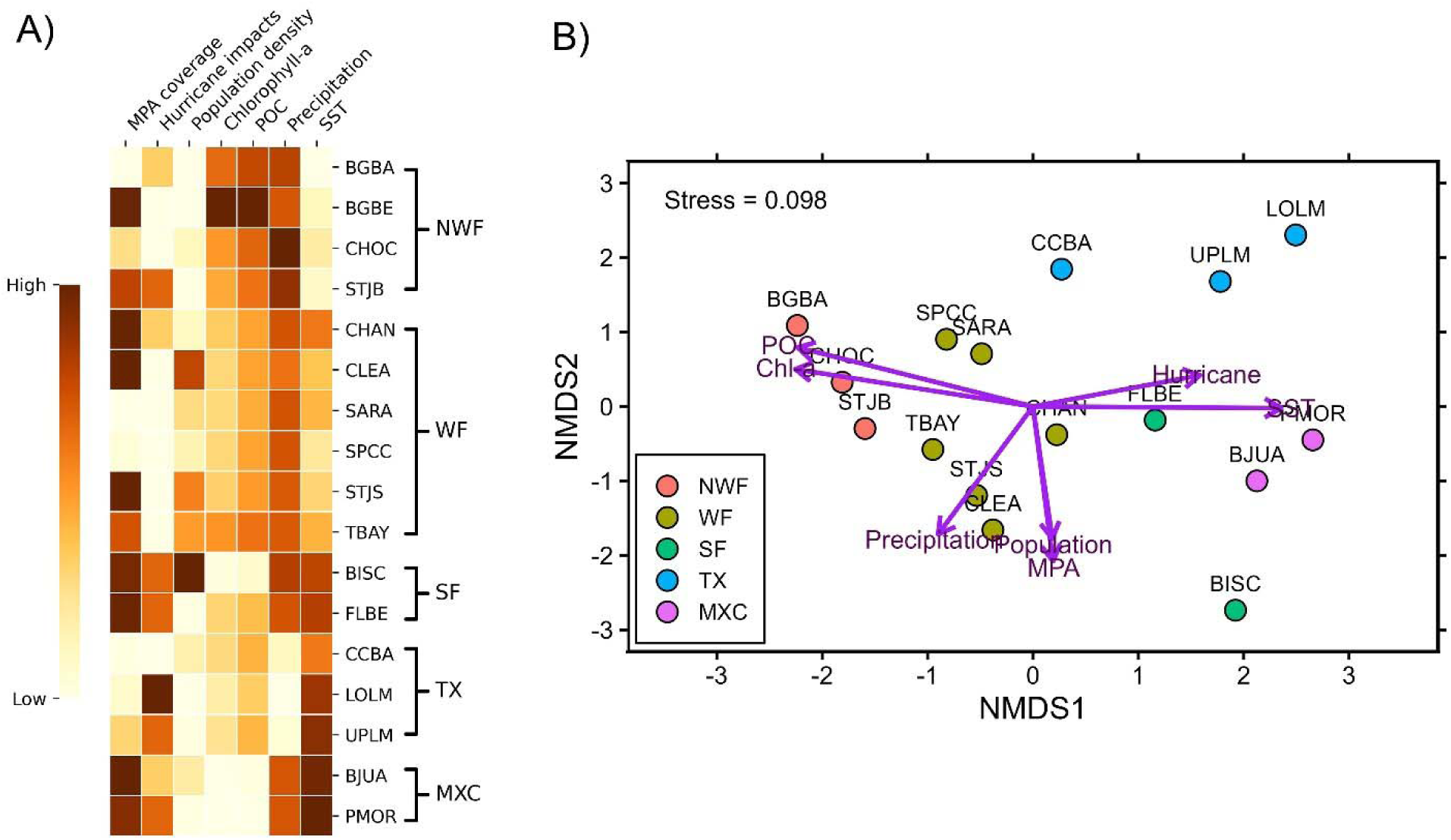
A) General characterization of seagrass sites with environmental variables of interest. Each variable follows a color gradient from the lowest to the highest average values across sites. B) NMDS ordination plot illustrating the similarities between seagrass sites and their gradients according to the characterization variables. See Table 1 for a description of site acronyms.

## 4. Discussion

### 4.1. Regional seagrass extent

The total seagrass areal extent quantified using satellite remote sensing in the Gulf of Mexico and Northwestern Caribbean, including areas of South and Southeast Florida such as the Florida Keys and Biscayne Bay was 19,405 km^2^. This is 44% lower than that estimated from the world seagrass distribution dataset (UNEP-WCMC and Short, 2021). That assessment reports 35,510 km^2^ of seagrass for this region, approximately, including areas that were not mapped in this study. The seagrass extent for the US Gulf of Mexico from 2017 reported by Handley and Lockwood (2020) was 6,683 km^2^; also higher in comparison to the 4,427 km^2^ from this study. Most of those distribution maps and extent estimates from the region are derived from detailed aerial photography and visual interpretations, supported by field monitoring (Handley et al., 2007; Yarbro and Carlson, 2016), limiting mapping in turbid and deeper areas. Gaps in reported seagrass extent still remain for areas such as the Florida Big Bend and Southwest Florida shelves, which apparently have the presence of extensive beds of *Halophila decipiens* up to 37 m depth and extent estimates of 12,000 km^2^ (Handley et al., 2007).

The differences in seagrass extent estimates among datasets are likely due to the limited capacity of satellite remote sensing to detect seagrass beds in optically visible waters (e.g., relatively clear and shallow waters) and pixel resolutions (Sentinel-2 at 10 m and Landsat at 30 m) that can produce these differences (Lizcano-Sandoval et al., 2022). Seagrasses in the region tend to be more abundant in shallow areas (<5 m depth) (Green and Short, 2003), but they can also grow in optically deeper waters (>10 m depth) from Cuba, the Yucatan peninsula, and the Southwest Florida shelf (de Almeida et al., 2022; Martínez-Daranas and Suárez, 2018; Palafox-Juárez and Liceaga-Correa, 2017) that were not possible to map with satellite imagery. Moreover, distribution datasets and monitoring programs may include both dense and sparse seagrass beds (sometimes referred as continuous and discontinuous seagrass), but we only consider that satellite data detects visible dense seagrass beds (usually >50% cover per pixel). Therefore, reported seagrass distributions are larger total extents than those reported by satellite mapping (Lizcano-Sandoval et al., 2022). Satellite and in situ monitoring data should ideally be used as complementary, and mapping methods should be standardized to allow more accurate comparisons across datasets (Neckles et al., 2012).

In total, we used 1,869 individual satellite images to run 2,304 classifications and map the seagrass distribution of the study region, including time series mapping. This allowed us to reconstruct time series of seagrass extent for 17 sites between 1987 and 2021. Although the seagrass mapping workflow used was semi-automated, the process required extensive user intervention for selecting images for mapping and post-processing (Lizcano-Sandoval et al., 2025, 2022). The approach led to seagrass classification accuracies >70 %, in general. Post-processing helped to reduce pixel noise and false positives in changes over time at a pixel level, given that classification outputs presented noises at some level. Seagrass mapping was challenging in most of the sites due to high spatial variability of water quality conditions, but collecting several images per year and segmenting and defining manageable site areas (e.g., Big Bend (A–E), Charlotte Harbor (North and South), Florida Bay (East and West), among others) helped to map seagrasses. This approach has been useful to enhance seagrass mapping at inter and intra-annual scales in sites such as Tampa Bay (Lizcano-Sandoval et al., 2025).

### 4.2. Seagrass extent trends and potential causes of change

Interpretations of changes in seagrass extent over time depend on the timeframe over which the trends are computed. For example, the trends we computed for seagrass extent in Lower and Upper Laguna Madre were -0.08% yr^-1^ and +0.18% yr^-1^, but the change between the initial and final year (i.e., 1987 vs. 2021) were 21% and -20%, respectively. Also, seagrass extent in 1987 for these two sites in Texas was different to the following years, probably as a response of the seagrass to some previous stressor that we were not able to identify. Our interpretations were based on changes in seagrass extent reported using the full interannual time series trends (or rates of change) derived from linear fitted models. Potential differences caused by different satellite sensors and resolutions are assumed to be small with our mapping approach. Lizcano-Sandoval et al. (2022) found that Landsat 8 typically underestimated seagrass extent by about 8% on average relative to maps derived with Sentinel-2. In multitemporal mapping it is challenging to quantify the potential error caused by resolution, but the use of Landsat and Sentinel-2 for detecting dense seagrass beds over time has shown high correlations with other temporal mapping methods using aerial imagery (Lizcano-Sandoval et al., 2025, 2022).

The highest rates of seagrass change were recorded in West Florida, specifically in St. Joseph Sound, Clearwater Harbor, and Tampa Bay. Sustained increase in seagrass extent in these sites is the result of improvements in water quality (e.g., reductions in total nitrogen and chlorophyll concentrations) since the 1980’s (Greening et al., 2014; Sherwood et al., 2017; Tomasko et al., 2018). Water quality in Tampa Bay improved as evidenced by an overall decrease in concentration of chlorophyll-a (-20%), nitrogen (-46%), phosphorus (-24%), and turbidity (-70%) between 1990–2020 (Lizcano-Sandoval et al., 2023). This overall trend was in spite of the discharge of 814 million liters of phosphate wastewater from the Piney Point plant into lower Tampa Bay in 2021 (Beck et al., 2022; Tomasko, 2025).

Sarasota Bay showed regular increases in seagrass extent year after year, but since 2018 there have been losses affecting the overall rate estimated over 1990–2021. These losses might have been intensified by the impact of Hurricane Irma in 2017 and the 2021 Piney Point spill (Beck et al., 2022; Tomasko, 2025). A high fraction of the West Florida sites showing significant increases in seagrass extent also were within MPA boundaries. The exception was Charlotte Harbor (North). Slight negative (and not significant) seagrass extent trends were observed in Charlotte Harbor (North) (-0.11% yr^-1^). Suspended particles, nutrients, and CDOM are known water quality drivers and possibly causes of seagrass change in Charlotte Harbor due to reduced light penetration (Brown et al., 2013; Corbett, 2006; Tomasko et al., 2005).

Choctawhatchee Bay also showed a significant increase in seagrass extent during the study period. This coincided with the large increases reported since 2007 (McDowell et al., 2018). However, seagrass extent in Choctawhatchee Bay is relatively small (∼5 km^2^) and showed high interannual variability (∼20%), suggesting that seagrass meadows in this site are susceptible to rapid extent changes. Furthermore, changes observed in Choctawhatchee Bay are potentially controlled by variations in light availability driven by water quality (influenced by storms, colored dissolved organic matter (CDOM), algae blooms), especially in the east side of the bay (Hoyer et al., 2015; McDowell et al., 2018; Yarbro and Carlson, 2016). Other sites in Northwest Florida such as Big Bend A and Big Bend C, located at the northwestern end of the Big Bend region in Florida, did not present significant changes in seagrass extent over time. Water quality has in the past impacted the seagrass habitats negatively in the Big Bend region (Yarbro et al., 2024). The Big Bend region frequently experiences events that alter the light quality for seagrasses, including algal blooms, resuspended sediments, and high CDOM (also known as dark water events) and nutrient discharge from the Ochlockonee, Steinhatchee, and Suwanee rivers (Xu et al., 2023; Yarbro et al., 2024).

In South Florida, the Florida Bay (East) and Biscayne Bay sites showed contrasting patterns in seagrass extent over time. Positive significant trends in seagrass extent were found for Florida Bay (East). Water quality conditions in Florida Bay are very complex. High salinity and temperature, sediment resuspension, high nitrogen and low phosphorous concentrations, recurrent algal blooms, and hypoxia all occur frequently in these shallow waters (Cannizzaro et al., 2019; de Fouw et al., 2024; Zhang et al., 2004). A massive seagrass die-off associated to hypersalinity events (>45 units) was reported for Florida Bay in 1987 (Robblee et al., 1991; Zieman et al., 1999), but seagrasses recovered after 1995 (Hall et al., 2021). Our estimates of an increase in seagrass extent in this area coincided with those ground observations. Biscayne Bay presented the steepest negative rate of change (-0.43% yr^-1^). Factors such as freshwater discharge controlling salinity variability and nutrient inputs, and habitat fragmentation, are reported as plausible causes of seagrass loss in Biscayne Bay (Lirman et al., 2014; Santos et al., 2016). Florida Bay and Biscayne Bay are surrounded by highly urbanized areas that historically have changed the freshwater flow from the Everglades and affected the water quality in those sites (Sklar et al., 2002; Steinman et al., 2002). Environmental damage to the Everglades water shed led to the development of the Comprehensive Everglades Restoration Program, one of the largest and most costly ecological restoration plans in history (Perry, 2008; Sklar et al., 2002).

In Texas, salinity and turbidity are potentially the main factors of seagrass change in the Laguna Madre system (Kowalski et al., 2018; Wilson and Dunton, 2018). However, propeller scarring by boats has been reported to have a negative impact in the seagrass habitats of south Texas (Dunton and Schonberg, 2002). The MPA coverage in these sites was <30%.

In the Mexican Caribbean, the two sites examined showed noticeably high SST, low chlorophyll-a and low POC concentrations, in comparison to the other seagrass sites in the Gulf. A gradual increase in nutrients due to groundwater discharge, along with coastal development has been observed in this region (Gómez et al., 2022; Null et al., 2014; van Tussenbroek, 2011). And, yet these sites show continuous growth in seagrass extent in what otherwise may be oligotrophic waters. A threat to seagrasses and corals that has emerged since 2011 are sargassum beaching events in this region of Mexico, as these can smother other marine life, diminish light availability, and lead to oxygen depletion when the sargasso algae die (Ávila-Mosqueda et al., 2025; Gómez et al., 2022; Martinez Ortiz et al., 2025).

Seagrasses of the Gulf of Mexico, Florida, and Mexican Caribbean are negatively affected by storms and hurricanes in the short-term, but they also show a high degree of resilience (Carlson et al., 2010; Tomasko et al., 2020; van Tussenbroek et al., 2014).

### 4.3. Seagrass conservation and management

Our results show that seagrass areal extent in the Gulf of Mexico, Florida, and Mexican Caribbean is in general stable and has increased in the last decade or more. However, sites such as Biscayne Bay, Charlotte Harbor (North), and Lower Laguna Madre may need management interventions and monitoring that benefit overall seagrass health and growth. Other sites not mapped temporally in our study require more frequent efforts to be monitored in the field and determine their change rates with higher accuracy. Spatial MPA coverage was not a clear indicator of resilience or seagrass health. Sites with good MPA coverage such as Biscayne Bay, Big Bend (East), and Charlotte Harbor (North) often showed negative change in seagrass extent or very small positive trends. However, in West Florida sites, high MPA coverage coincided with positive changes in seagrass extent. The establishment of national estuary programs in West Florida, along with legislation and regulation of wastewater and stormwater discharges have resulted in an increase of seagrass extent (Greening and Janicki, 2006; Sherwood et al., 2016; Tomasko et al., 2018). The temporal dynamics of seagrasses in the Mexican Caribbean seem to reflect a healthy ecosystem. However, if eutrophication continues an ecosystem balance disruption might occur that also depends on the coral reefs and other organisms that inhabit these coasts.

There is ample evidence that effective legislation and management of nutrient inputs help improve the extent of seagrass beds by improving water quality and allowing more light to reach the seagrass habitat (Beck et al., 2018; Choice et al., 2014). Tampa Bay is a case of success in restoring water quality that positively impacted seagrass habitats (Sherwood et al., 2016; Tomasko et al., 2018). Also, long-term active restoration efforts have shown to be positive for seagrasses in the US east coast (Orth et al., 2017). Increasing efforts on managing and conserving seagrass habitats and ecosystems that surround them (e.g., coral reefs, mangroves, salt marshes) would further enhance their overall health state in a region under constant pressures from multiple stressors related to coastal development and climate change.

## 5. Conclusions

Overall, the seagrass habitats in the Gulf of Mexico and Northwestern Caribbean have been stable or have increased their areal extent during 1987–2021. Long-term changes were best characterized by considering annual data and the interannual variations in areal extent, and not simply by comparing the initial and final years (year_f_ – year_i_) as indicators of seagrass change. Further, seagrass meadows are dynamic habitats that can expand or contract from year to year due to different causes. Positive seagrass changes observed in West Florida are an example of management approaches that achieve the desired positive goal for seagrass conservation and recovery, that could be replicated to other sites in the Gulf of Mexico and Caribbean.

We demonstrated that satellite remote sensing and cloud-based geospatial platforms open the possibility for large-scale, intensive seagrass areal extent monitoring by anyone. This access benefits conservation and management of seagrass and other coastal habitats.

There is still work needed to improve seagrass mapping methods. Substantial work is required to validate products, in automation of classifications and deriving assessments. Availability of georeferenced seagrass presence records is a necessity to achieve ongoing mapping across seagrass habitats to reduce bias, effort, and processing times. Temporally resolved maps of seagrasses aid in detecting change, prioritizing resources for ecological and biogeochemical studies, and providing actionable information to decision-makers and policy.

## Acknowledgements

This work was made possible by support provided to L. Lizcano-Sandoval by the Fulbright-Colciencias Program of Colombia, the College of Marine Science at the University of South Florida (Norman Blake Endowed Memorial Fellowship, Sanibel-Captiva Shell Club/Mary & Al Bridell Memorial Fellowship), and the NASA Future Investigators in NASA Earth and Space Science and Technology (FINESST) scholarship. Support was provided by the Marine Biodiversity Observation Network (MBON: NASA grants NNX14AP62A; 80NSSC20K0017; 80NSSC22K1779; NOAA IOOS grant NA19NOS0120199), the Gulf of America Coastal Ocean Observing System (GCOOS/IOOS Cooperative Agreement NA16NOS0120018), and the Tampa Bay Estuary Program (TBEP: 2500-1888). We thank all the organizations that made seagrass monitoring data and maps available to the public.

