## Supplemental Information for "Seagrass Extent Expansion in the Gulf of Mexico and Northwestern Caribbean (1987–2021)"

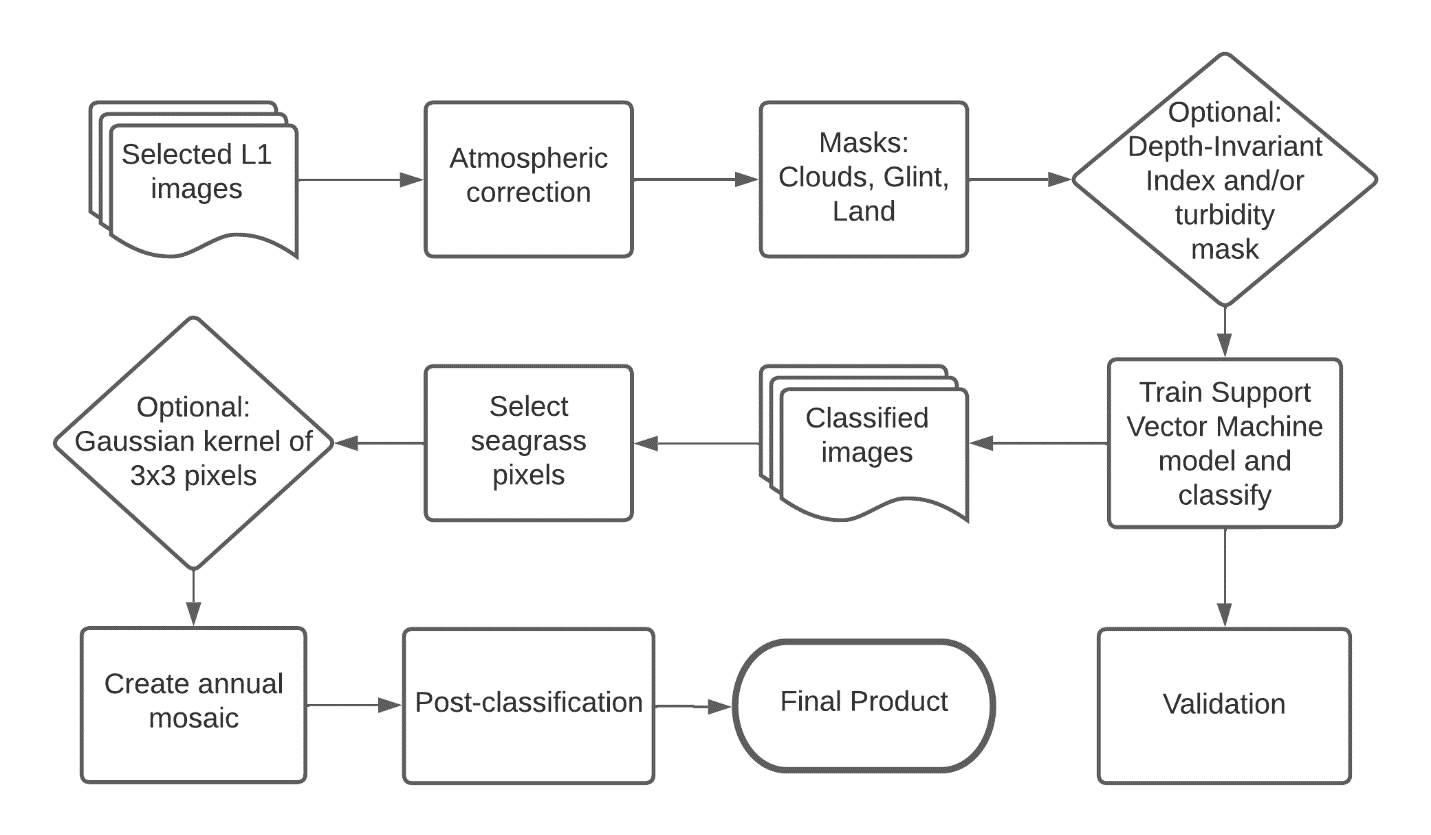

**Figure S1.** Seagrass mapping workflow described by Lizcano-Sandoval et al. (2022) with two optional steps added (diamond boxes).

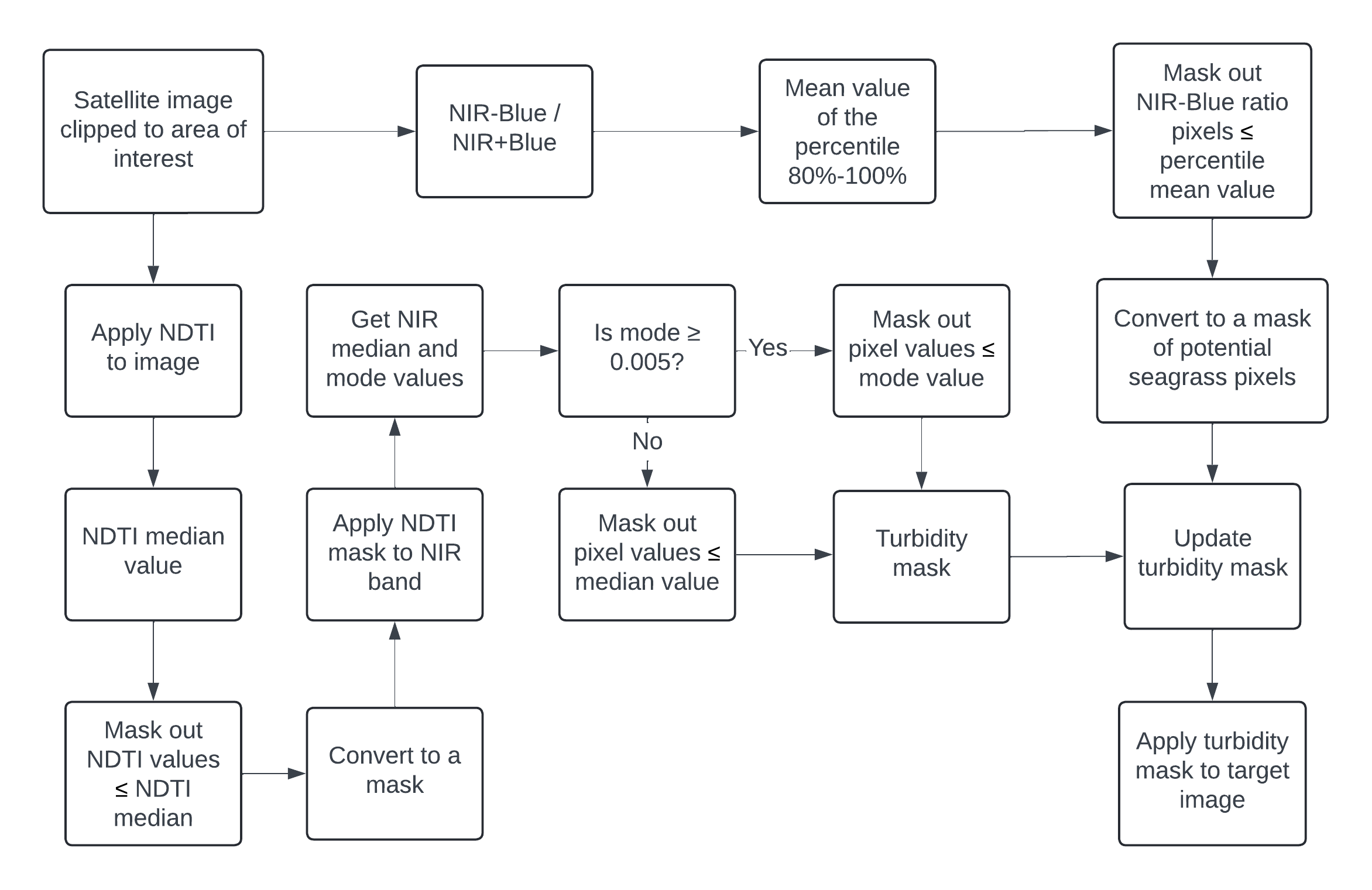

**Figure S2.** Turbidity masking process in waters with seagrass beds presence.

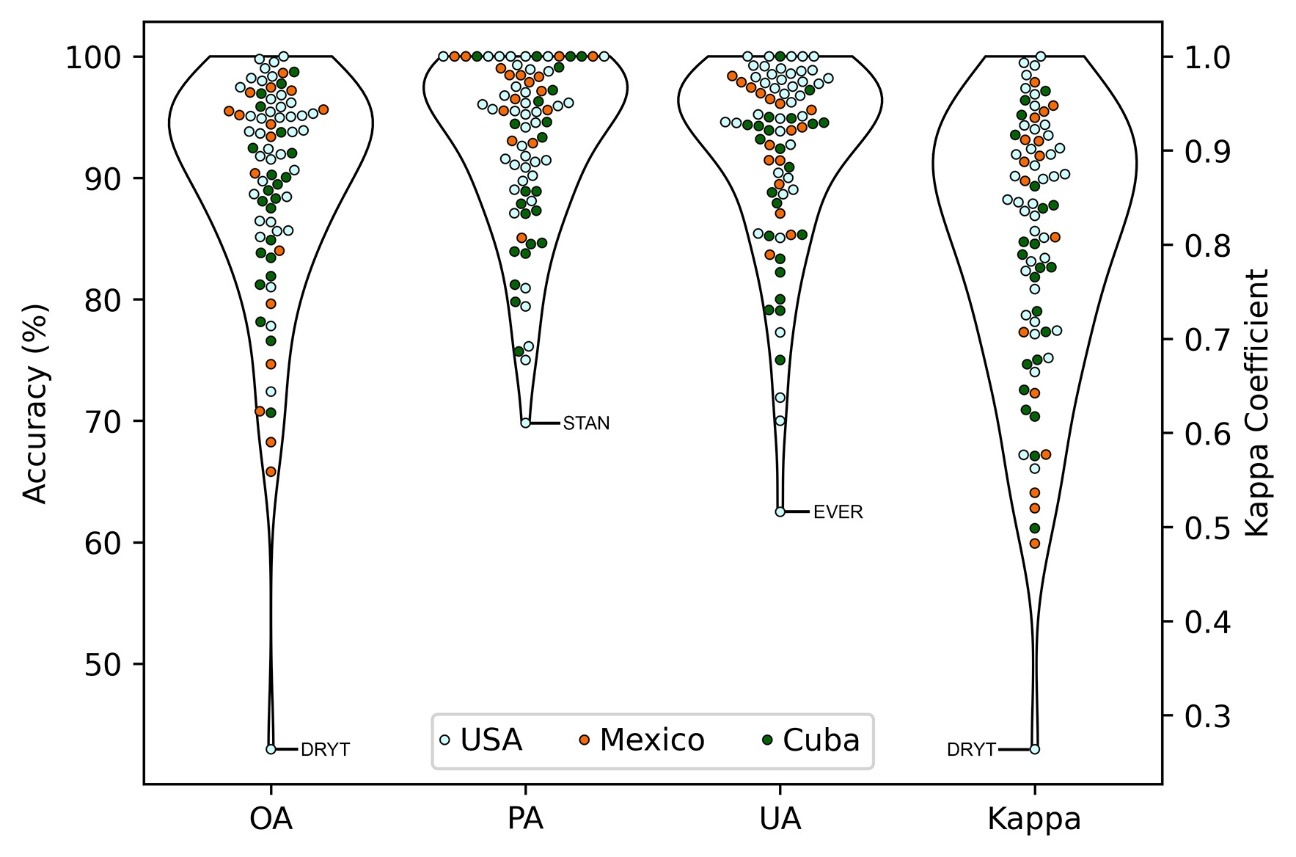

**Figure S3.** Average seagrass regional distribution accuracies grouped by country. The site with the respective lowest accuracy or Kappa coefficient is indicated. OA: overall accuracy; PA: seagrass producer’s accuracy; UA: seagrass user’s accuracy.

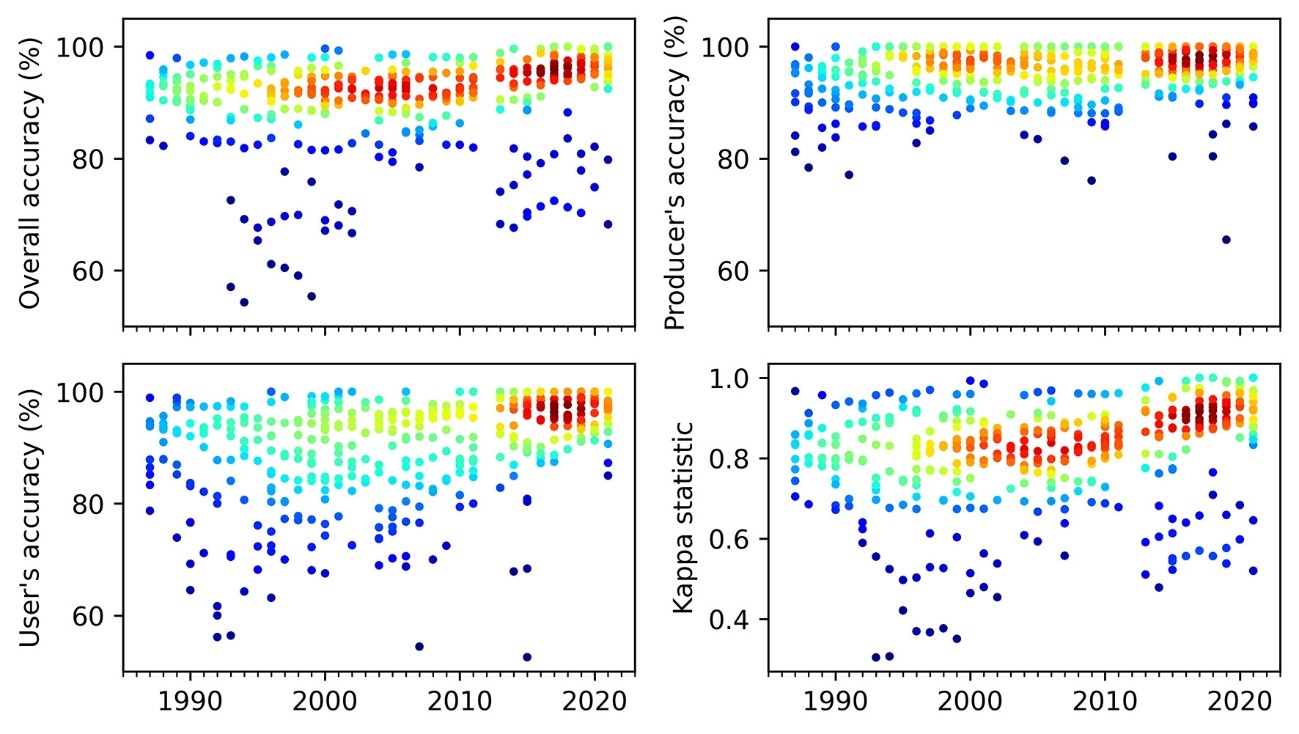

**Figure S4.** Average validation accuracies of annual seagrass classifications over 1987–2021. Blue indicates few dots density and red indicates high dots density.

**Table S1.** Seagrass extent of the study areas in the Gulf of Mexico and Northern Caribbean. Coordinates represent the centroid of each area. Sites selected for temporal mapping are also indicated as “Yes+TS”.

| **Country** | **State** | **Site** | **Abbr.** | **Lat** | **Lon** | **Mapped** | **Year** | **n images** | **Seagrass extent (km^2^)** |
| --- | --- | --- | --- | --- | --- | --- | --- | --- | --- |
| USA | Alabama | Mobile Bay | MOBI | -87.96 | 30.42 | No |  |  |  |
| USA | Florida | Big Bend A | BGBA | -84.21 | 29.90 | Yes+TS | 2020 | 2 | 30.91 |
| USA | Florida | Big Bend B | BGBB | -84.27 | 30.03 | Yes | 2019 | 4 | 39.54 |
| USA | Florida | Big Bend C | BGBC | -84.09 | 30.06 | Yes | 2019 | 11 | 97.06 |
| USA | Florida | Big Bend D | BGBD | -83.85 | 29.93 | Yes | 2019 | 9 | 466.69 |
| USA | Florida | Big Bend E | BGBE | -83.58 | 29.72 | Yes+TS | 2021 | 7 | 113.29 |
| USA | Florida | Big Bend F | BGBF | -83.40 | 29.50 | Yes | 2019 | 4 | 107.97 |
| USA | Florida | Biscayne Bay | BISC | -80.19 | 25.56 | Yes+TS | 2021 | 6 | 456.85 |
| USA | Florida | Cedar Key | CEDA | -83.08 | 29.18 | Yes | 2019 | 10 | 20.58 |
| USA | Florida | Charlotte Harbor (North) | CHAN | -82.21 | 26.86 | Yes+TS | 2021 | 6 | 41.92 |
| USA | Florida | Charlotte Harbor (South) | CHAS | -82.13 | 26.56 | Yes | 2019 | 5 | 102.80 |
| USA | Florida | Choctawhatchee | CHOC | -86.45 | 30.43 | Yes+TS | 2021 | 8 | 5.41 |
| USA | Florida | Clearwater Harbor | CLEA | -82.79 | 27.96 | Yes+TS | 2021 | 2 | 9.75 |
| USA | Florida | Dry Tortugas | DRYT | -83.86 | 24.66 | Yes | 2019 | 5 | 28.69 |
| USA | Florida | Estero Bay | ESTE | -81.89 | 26.43 | Yes | 2019 | 5 | 2.29 |
| USA | Florida | Everglades | EVER | -81.29 | 25.62 | Yes | 2019 | 3 | 3.06 |
| USA | Florida | Florida Bay (East) | FLBE | -80.76 | 25.06 | Yes+TS | 2021 | 12 | 982.50 |
| USA | Florida | Florida Bay (West) | FLBW | -81.25 | 24.96 | Yes | 2019 | 6 | 465.45 |
| USA | Florida | Franklin County | FRAN | -84.69 | 29.79 | Yes | 2019 | 12 | 22.69 |
| USA | Florida | Indian River Lagoon (North) | IRLN | -80.27 | 27.39 | No |  |  |  |
| USA | Florida | Indian River Lagoon (South) | IRLS | -80.67 | 28.46 | No |  |  |  |
| USA | Florida | Lower FL Keys | LFLK | -81.56 | 24.63 | Yes | 2019 | 9 | 1,426.27 |
| USA | Florida | Marquesas Keys | MARQ | -82.28 | 24.56 | Yes | 2019 | 4 | 181.96 |
| USA | Florida | Middle FL Keys | MFLK | -80.88 | 24.79 | Yes | 2019 | 9 | 652.88 |
| USA | Florida | Pensacola Bay | PENS | -87.07 | 30.36 | Yes | 2019 | 9 | 9.62 |
| USA | Florida | Perdido Bay | PERD | -87.50 | 30.29 | Yes | 2019 | 5 | 1.31 |
| USA | Florida | Rookery Bay A | ROOK | -81.69 | 25.90 | Yes | 2019 | 4 | 1.25 |
| USA | Florida | Rookery Bay B | ROOB | -81.53 | 25.85 | No |  |  |  |
| USA | Florida | Sarasota Bay | SARA | -82.53 | 27.34 | Yes+TS | 2021 | 2 | 25.79 |
| USA | Florida | Springs Coast A | SPCA | -82.83 | 28.82 | Yes | 2019 | 11 | 338.72 |
| USA | Florida | Springs Coast B | SPCB | -82.81 | 28.57 | Yes | 2019 | 9 | 465.28 |
| USA | Florida | Springs Coast C | SPCC | -82.82 | 28.32 | Yes+TS | 2021 | 8 | 299.63 |
| USA | Florida | St. Andrew Bay | STAN | -85.68 | 30.15 | Yes | 2019 | 3 | 34.72 |
| USA | Florida | St. Joseph Bay | STJB | -85.35 | 29.78 | Yes+TS | 2021 | 13 | 24.46 |
| USA | Florida | St. Joseph Sound | STJS | -82.79 | 28.11 | Yes+TS | 2021 | 4 | 42.38 |
| USA | Florida | Tampa Bay | TBAY | -82.59 | 27.77 | Yes+TS | 2021 | 24 | 82.14 |

**Table S1.** Continued

| **Country** | **State** | **Site** | **Abbr.** | **Lat** | **Lon** | **Mapped** | **Year** | **n images** | **Seagrass extent (km^2^)** |
| --- | --- | --- | --- | --- | --- | --- | --- | --- | --- |
| USA | Florida | Upper FL Keys | UFLK | -80.36 | 25.12 | Yes | 2019 | 11 | 501.26 |
| USA | Louisiana | Chandeleur Island | CHAI | -88.91 | 29.82 |  | 2020 | 5 | 12.96 |
| USA | Mississippi | Mississippi Sound | MISS | -88.64 | 30.27 | No |  |  |  |
| USA | Texas | Aransas Bay | ARAN | -96.85 | 28.20 | Yes | 2020 | 15 | 71.22 |
| USA | Texas | Corpus Christi Bay | CCBA | -97.28 | 27.80 | Yes+TS | 2021 | 7 | 28.22 |
| USA | Texas | Galveston Bay | GALV | -94.87 | 29.45 | Yes | 2020 | 7 | 1.14 |
| USA | Texas | Lower Laguna Madre | LOLM | -97.34 | 26.44 | Yes+TS | 2021 | 11 | 291.84 |
| USA | Texas | Matagorda Bay | MATB | -96.33 | 28.61 | Yes | 2020 | 6 | 7.95 |
| USA | Texas | Upper Laguna Madre | UPLM | -97.44 | 27.25 | Yes+TS | 2021 | 10 | 176.17 |
| Mexico | Campeche | Champotón | CHAM | -90.79 | 19.36 | Yes | 2020 | 8 | 112.52 |
| Mexico | Campeche | Laguna de Términos | LTER | -91.53 | 18.61 | Yes | 2020 | 4 | 82.04 |
| Mexico | Campeche | Petenes | PETE | -90.58 | 20.29 | Yes | 2020 | 4 | 1,328.31 |
| Mexico | Tamaulipas | Laguna Madre (Mexico) | LMMX | -97.64 | 25.01 | Yes | 2020 | 4 | 495.35 |
| Mexico | Tamaulipas | Laguna Morales | LMOR | -97.76 | 23.70 | Yes | 2020 | 10 | 10.20 |
| Mexico | Tamaulipas | Laguna San Andres | LSAN | -97.86 | 22.67 | Yes | 2020 | 4 | 16.68 |
| Mexico | Veracruz | Laguna Tamiahua | LTAM | -97.58 | 21.64 | Yes | 2020 | 6 | 23.56 |
| Mexico | Veracruz | Veracruz | VERA | -95.87 | 19.10 | No |  |  |  |
| Mexico | Quintana Roo | Laguna Yalahau | YALA | -87.28 | 21.49 | Yes | 2020 | 12 | 108.91 |
| Mexico | Yucatan | Yucatan Coast | YUCA | -89.05 | 21.41 | Yes | 2020 | 3 | 370.15 |
| Mexico | Quintana Roo | Akumal Coast | AKUM | -87.33 | 20.17 | Yes | 2020 | 5 | 10.11 |
| Mexico | Quintana Roo | Banco Chinchorro | BCHI | -87.34 | 18.59 | Yes | 2020 | 7 | 13.87 |
| Mexico | Quintana Roo | Benito Juarez | BJUA | -86.79 | 21.01 | Yes+TS | 2021 | 6 | 14.99 |
| Mexico | Quintana Roo | Carrillo Puerto | CARP | -87.54 | 19.48 | Yes | 2020 | 4 | 38.01 |
| Mexico | Quintana Roo | Catoche Cape | CATO | -86.97 | 21.49 | Yes | 2020 | 7 | 218.27 |
| Mexico | Quintana Roo | Cozumel Island | COZI | -86.88 | 20.46 | No |  |  |  |
| Mexico | Quintana Roo | Puerto Blanco | PBLA | -87.96 | 18.56 | Yes | 2020 | 4 | 13.62 |
| Mexico | Quintana Roo | Puerto Morelos | PMOR | -86.87 | 20.85 | Yes+TS | 2021 | 12 | 6.71 |
| Cuba | Camagüey | Archipelago Camagüey (East) | CAME | -77.67 | 22.07 | Yes | 2021 | 26 | 36.34 |
| Cuba | Camagüey | La Gloria Bay | GLOR | -77.61 | 21.81 | Yes | 2021 | 5 | 100.31 |
| Cuba | Camagüey | Jardines de la Reina | JARD | -79.00 | 20.95 | Yes | 2021 | 14 | 784.04 |
| Cuba | Camagüey | Jiguey Bay | JIGU | -78.15 | 22.18 | No |  |  |  |
| Cuba | Camagüey | Nuevitas Bay | NUEV | -77.24 | 21.54 | Yes | 2021 | 8 | 52.78 |
| Cuba | Camagüey | Santa Lucia | STLU | -77.02 | 21.55 | No |  |  |  |

**Table S1.** Continued

| **Country** | | **State** | **Site** | **Abbr.** | **Lat** | **Lon** | **Mapped** | **Year** | **n images** | **Seagrass extent (km^2^)** |
| --- | --- | --- | --- | --- | --- | --- | --- | --- | --- | --- |
|  | Cuba | Ciego de Ávila | Archipelago Camagüey (West) | CAMW | -78.48 | 22.51 | Yes | 2021 | 26 | 96.20 |
|  | Cuba | Ciego de Ávila | Perros Bay | PERR | -78.55 | 22.39 | No |  |  |  |
|  | Cuba | Cienfuegos | Cienfuegos Bay | CIEN | -80.47 | 22.12 | Yes | 2021 | 10 | 0.85 |
|  | Cuba | Granma | Angostura | ANGO | -77.24 | 20.61 | Yes | 2021 | 5 | 15.03 |
|  | Cuba | Granma | Cruz Cape | CRUZ | -77.72 | 19.92 | Yes | 2021 | 4 | 13.27 |
|  | Cuba | Granma | Manzanillo | MANZ | -77.12 | 20.42 | No |  |  |  |
|  | Cuba | Guantanamo | Guantanamo Bay | GUAN | -75.15 | 19.93 | No |  |  |  |
|  | Cuba | Holguin | Moa | MOAC | -74.90 | 20.67 | No |  |  |  |
|  | Cuba | Holguin | Nipe Bay | NIPE | -75.63 | 20.77 | No |  |  |  |
|  | Cuba | Isla de la Juventud | Batabanó Gulf (East) | BATE | -82.22 | 22.10 | Yes | 2021 | 4 | 1,360.98 |
|  | Cuba | Isla de la Juventud | Batabanó Gulf (West) | BATW | -83.02 | 21.95 | Yes | 2021 | 12 | 3,107.97 |
|  | Cuba | Isla de la Juventud | Archipelago Canarreos | CANA | -81.41 | 21.67 | Yes | 2021 | 5 | 76.93 |
|  | Cuba | Isla de la Juventud | Diego Perez Cays | DIEG | -81.43 | 21.91 | Yes | 2021 | 4 | 97.65 |
|  | Cuba | Isla de la Juventud | Ciénaga de Zapata (North) | ZAPN | -81.91 | 22.55 | No |  |  |  |
|  | Cuba | Isla de la Juventud | Ciénaga de Zapata (South) | ZAPS | -81.90 | 22.21 | Yes | 2021 | 8 | 371.18 |
|  | Cuba | Matanzas | Blancos Cays | BLAN | -81.31 | 22.10 | Yes | 2021 | 10 | 85.30 |
|  | Cuba | Matanzas | Cardenas Bay | CARD | -81.01 | 23.17 | Yes | 2021 | 4 | 222.05 |
|  | Cuba | Pinar del Río | Archipelago Los Colorados | COLO | -83.84 | 22.73 | Yes | 2021 | 9 | 331.09 |
|  | Cuba | Pinar del Río | Guanahacabibes Gulf | GUAG | -84.59 | 22.12 | Yes | 2021 | 10 | 661.37 |
|  | Cuba | Sancti Spíritus | Buena Vista Bay | BVIS | -79.03 | 22.47 | Yes | 2021 | 20 | 562.98 |
|  | Cuba | Sancti Spíritus | Trinidad | TRIN | -79.88 | 21.69 | Yes | 2021 | 12 | 31.83 |
|  | Cuba | Villa Clara | Carahatas Bay | CARA | -80.49 | 23.09 | Yes | 2021 | 13 | 320.91 |
|  | Cuba | Villa Clara | Novillo Bay | NOVI | -79.98 | 22.96 | Yes | 2021 | 6 | 158.34 |
|  | Cuba | Villa Clara | San Juan Bay | SJUA | -79.54 | 22.70 | Yes | 2021 | 5 | 380.13 |

**Table S2.** Characteristics of satellite sensors and relevant bands considered for seagrass mapping.

| **Satellite Sensor** | **Pixel Size (m)** | **Bands of Interest (nm)** | **Band Names** | **Period Covered in this Study** | **Revisit Time (Days)** |
| --- | --- | --- | --- | --- | --- |
| Landsat-5 TM | 30 | 485, 560, 660 | Blue, Green, Red | 1987–2012 | 16 |
| Landsat-7 ETM+ | 30 | 485, 560, 660 | Blue, Green, Red | 1999–2002 | 16 |
| Landsat-8 OLI | 30 | 443, 482, 561, 655 | Coastal, Blue, Green, Red | 2013–2016 | 16 |
| Sentinel-2A/B MSI | 10 | 443*, 482, 561, 655 | Coastal, Blue, Green, Red | 2016–2021 | ~5 |

*Native resolution at 60 m resolution, but reprojected to 10 m.

**Table S3.** Sources of seagrass distribution and ground-truth data.

| #N | Country | State | Location | Year | Author(s) | Title | Type | Publisher | Vol: pp | URL/doi |
| --- | --- | --- | --- | --- | --- | --- | --- | --- | --- | --- |
| 1 | USA | Florida | Central West Florida | 1982-2020 | Southwest Florida Water Management District | Aerial seagrass mapping data | Dataset | Southwest Florida Water Management District | NA | [Website](https://data-swfwmd.opendata.arcgis.com) |
| 2 | USA | Florida | Florida | 2016 | Yarbro, L.A., Carlson, P.R.J. | Seagrass integrated mapping and monitoring program: Mapping and monitoring report No. 2 | Report | Fish and Wildlife Research Institute | NA | [Website](https://myfwc.com/research/habitat/seagrasses/projects/active/simm/simm-reports/) |
| 3 | USA | Florida | Springs Coast | 2016 | Baumstark et al. | Mapping seagrass and colonized hard bottom in Springs Coast, Florida using WorldView-2 satellite imagery | Journal Article | Estuar. Coast. Shelf Sci. | 181: 83–92 | [10.1016/j.ecss.2016.08.019](https://doi.org/10.1016/j.ecss.2016.08.019%20) |
| 4 | USA | Florida | St. Joseph Bay | 2014 | Hill et al. | Evaluating light availability, seagrass biomass, and productivity using hyperspectral airborne remote sensing in Saint Joseph's Bay, Florida | Journal Article | Estuaries and Coasts | 37: 1467–1489 | [10.1007/s12237-013-9764-3](https://doi.org/10.1007/s12237-013-9764-3) |
| 5 | USA | Florida | St. Joseph Bay | 2022 | Lebrasse et al. | Temporal stability of seagrass extent, leaf area, and carbon storage in St. Joseph Bay, Florida: a semi-automated remote sensing analysis | Journal Article | Estuaries and Coasts | NA | [10.1007/s12237-022-01050-4](https://doi.org/10.1007/s12237-022-01050-4) |
| 6 | USA | Florida | St. Joseph Bay, Tampa Bay, St. George Sound | 2020 | Coffer et al. | Performance across WorldView-2 and RapidEye for reproducible seagrass mapping | Journal Article | Remote Sens. Environ. | 250: 112036 | [10.1016/j.rse.2020.112036](https://doi.org/10.1016/j.rse.2020.112036) |
| 7 | USA | Florida | St. Joseph Sound, Clearwater Harbor | 2012 | Meyer et al. | Seagrass resource assessment using remote sensing methods in St. Joseph Sound and Clearwater Harbor, Florida, USA | Journal Article | Environ. Monit. Assess. | 184: 1131–1143 | [10.1007/s10661-011-2028-4](https://doi.org/10.1007/s10661-011-2028-4) |
| 8 | USA | Florida | St. Joseph Sound, Clearwater Harbor | 2012 | Pu et al. | Discrimination of seagrass species and cover classes with in situ hyperspectral data | Journal Article | J. Coast. Res. | 285: 1330–1344 | [10.2112/jcoastres-d-11-00229.1](https://doi.org/10.2112/jcoastres-d-11-00229.1) |
| 9 | USA | Florida | St. Joseph Sound, Clearwater Harbor | 2012 | Pu et al. | Mapping and assessing seagrass along the western coast of Florida using Landsat TM and EO-1 ALI/Hyperion imagery | Journal Article | Estuar. Coast. Shelf Sci. | 115: 234–245 | [10.1016/j.ecss.2012.09.006](https://doi.org/10.1016/j.ecss.2012.09.006) |
| 10 | USA | Florida | St. Joseph Sound, Clearwater Harbor | 2013 | Pu, R., Bell, S. | A protocol for improving mapping and assessing of seagrass abundance along the West Central Coast of Florida using Landsat TM and EO-1 ALI/Hyperion images | Journal Article | ISPRS J. Photogramm. Remote Sens. | 83: 116–129 | [10.1016/j.isprsjprs.2013.06.008](https://doi.org/10.1016/j.isprsjprs.2013.06.008) |
| 11 | USA | Florida | St. Joseph Sound, Clearwater Habor | 2014 | Pu et al. | Mapping and assessing seagrass bed changes in Central Florida’s west coast using multitemporal Landsat TM imagery | Journal Article | Estuar. Coast. Shelf Sci. | 149: 68–79 | [10.1016/j.ecss.2014.07.014](https://doi.org/10.1016/j.ecss.2014.07.014) |

**Table S3.** Continued

| #N | Country | State | Location | Year | Author(s) | Title | Type | Publisher | Vol: pp | URL/doi |
| --- | --- | --- | --- | --- | --- | --- | --- | --- | --- | --- |
| 12 | USA | Florida | St. Joseph Sound, Clearwater Harbor | 2017 | Pu, R., Bell, S. | Mapping seagrass coverage and spatial patterns with high spatial resolution IKONOS imagery | Journal Article | Int. J. Appl. Earth Obs. Geoinf. | 54: 145–158 | [10.1016/j.jag.2016.09.011](https://doi.org/10.1016/j.jag.2016.09.011) |
| 13 | USA | Florida | Tampa Bay | 2017 | Sherwood et al. | Tampa Bay (Florida, USA): Documenting seagrass recovery since the 1980’s and reviewing the benefits | Journal Article | Southeast. Geogr. | 57: 294–319 | [10.1353/sgo.2017.0026](https://doi.org/10.1353/sgo.2017.0026) |
| 14 | USA | Florida | Tampa Bay | 1998-2020 | Tampa Bay Estuary Program | In situ seagrass monitoring data | Dataset | Tampa Bay Estuary Program | NA | [Website](https://tbep.org/seagrass-assessment/) |
| 15 | USA | Florida | West Florida | 1986 | Iverson, R.L., Bittaker, H.F. | Seagrass distribution and abundance in eastern Gulf of Mexico coastal waters | Journal Article | Estuar. Coast. Shelf Sci. | 22: 577–602 | 10.1016/0272-7714(86)90015-6 |
| 16 | USA | Florida, Texas | Pensacola Bay, Galveston Bay | 2021 | zu Ermgassen et al. | Estimating and applying fish and invertebrate density and production enhancement from seagrass, salt marsh edge, and oyster reef nursery habitats in the Gulf of Mexico | Journal Article | Estuaries and Coasts | 44: 1588–1603 | [10.1007/s12237-021-00935-0](https://doi.org/10.1007/s12237-021-00935-0) |
| 17 | USA | Louisiana | Chandeleur Islands | 2017 | Darnell et al. | Spatial and Temporal Patterns in *Thalassia testudinum* Leaf Tissue Nutrients at the Chandeleur Islands, Louisiana, USA | Journal Article | Estuaries and Coasts | 40: 1288–1300 | [10.1007/s12237-017-0229-y](https://doi.org/10.1007/s12237-017-0229-y) |
| 18 | USA | Louisiana | Chandeleur Islands | 2017 | Kenworthy et al. | Seagrass response following exposure to Deepwater Horizon oil in the Chandeleur Islands, Louisiana (USA) | Journal Article | Mar. Ecol. Prog. Ser. | 576: 145–161 | [10.3354/meps11983](https://doi.org/10.3354/meps11983) |
| 19 | USA | Mississippi | Horn Island | 2010 | Lucas, K., Carter, G.A. | Decadal Changes in Habitat-Type Coverage on Horn Island, Mississippi, U.S.A. | Journal Article | J. Coast. Res. | 26: 1142-1148 | [10.2112/JCOASTRES-D-09-00018.1](https://doi.org/10.2112/JCOASTRES-D-09-00018.1) |
| 20 | USA | Mississippi | Horn Island | 2013 | Lucas, K., Carter, G.A. | Change in distribution and composition of vegetated habitats on Horn Island, Mississippi, northern Gulf of Mexico, in the initial five years following Hurricane Katrina | Journal Article | Geomorphology | 199: 129-137 | 10.1016/j.geomorph.2012.11.010 |
| 21 | USA | Mississippi | Mississippi barrier islands | 2011 | Carter et al. | Historical changes in seagrass coverage on the Mississippi barrier islands, northern Gulf of Mexico, determined from vertical aerial imagery (1940–2007) | Journal Article | Geocarto Int. | 26: 663-673 | 10.1080/10106049.2011.620634 |
| 22 | USA | Mississippi, Louisiana | Mississippi and Chandeleur Sounds | 2014 | Pham et al. | Seagrasses in the Mississippi and Chandeleur Sounds and problems associated with decadal-scale change detection | Journal Article | Gulf of Mexico Science | 32: 24-43 | [10.18785/goms.3201.03](https://doi.org/10.18785/goms.3201.03) |
| 23 | USA | Texas | Galveston Bay | 1996 | Galveston Bay Estuary Program | Galveston Bay Status and Trends | Dataset | Galveston Bay Estuary Program | NA | [Website](https://www.galvbaydata.org/) |

**Table S3.** Continued

| #N | Country | State | Location | Year | Author(s) | Title | Type | Publisher | Vol: pp | URL/doi |
| --- | --- | --- | --- | --- | --- | --- | --- | --- | --- | --- |
| 24 | USA | Texas | Galveston Bay; Matagorda Bay | 2015 | Texas Parks & Wildlife | Aerial seagrass mapping data | Dataset | Texas Parks & Wildlife | NA | [Website](https://tpwd.texas.gov/landwater/water/habitats/coastal-fisheries-habitat-assessment-team/) |
| 25 | USA | Texas | Laguna Madre | 1993 | Quammen, M.L., Onuf, C.P. | Laguna madre: Seagrass changes continue decades after salinity reduction | Journal Article | Estuaries | 16: 302–310 | [10.2307/1352503](https://doi.org/10.2307/1352503) |
| 26 | USA | Texas | Laguna Madre | 1996 | Onuf, C.P. | Biomass patterns in seagrass meadows of the Laguna Madre, Texas | Journal Article | Bull. Mar. Sci. | 58: 404–420 | NA |
| 27 | USA | Texas | Laguna Madre | 2002 | Dunton, K.H., Schonberg, S.V. | Assessment of propeller scarring in seagrass beds of the South Texas Coast | Journal Article | J. Coast. Res. | 37: 100–110 | NA |
| 28 | USA | Texas | Laguna Madre | 2017 | Larkin et al. | A map-based approach to assessing genetic diversity, structure, and connectivity in the seagrass *Halodule wrightii* | Journal Article | Mar. Ecol. Prog. Ser. | 567: 95–107 | [10.3354/meps12037](https://doi.org/10.3354/meps12037) |
| 29 | USA | Texas | Laguna Madre | 2018 | Kowalski et al. | A Comparison of Salinity Effects from Hurricanes Dolly (2008) and Alex (2010) in a Texas Lagoon System | Journal Article | J. Coast. Res. | 34: 1429–1438 | [10.2112/JCOASTRES-D-18-00011.1](https://doi.org/10.2112/JCOASTRES-D-18-00011.1) |
| 30 | USA | Texas | Laguna Madre | 2018 | Wilson, S.S., Dunton, K.H. | Hypersalinity during regional drought drives mass mortality of the seagrass *Syringodium filiforme* in a subtropical lagoon | Journal Article | Estuaries and Coasts | 41: 855–865 | [10.1007/s12237-017-0319-x](https://doi.org/10.1007/s12237-017-0319-x) |
| 31 | USA | Texas | Laguna Madre | 2019 | Su, L., Huang, Y. | Seagrass resource assessment using Worldview-2 imagery in the Redfish Bay, Texas | Journal Article | J. Mar. Sci. Eng. | 7: 98 | [10.3390/jmse7040098](https://doi.org/10.3390/jmse7040098) |
| 32 | USA | Texas | Laguna Madre | 2020 | Bittner et al. | Using species distribution models to guide seagrass management | Journal Article | Estuar. Coast. Shelf Sci. | 240: 106790 | [10.1016/j.ecss.2020.106790](https://doi.org/10.1016/j.ecss.2020.106790) |
| 33 | USA | Texas | Laguna Madre, Corpus Christi Bay | 2017 | Congdon et al. | Evaluation of relationships between cover estimates and biomass in subtropical seagrass meadows and application to landscape estimates of carbon storage | Journal Article | Southeast. Geogr. | 57: 231–245 | [10.1353/sgo.2017.0023](https://doi.org/10.1353/sgo.2017.0023) |
| 34 | USA | Texas | Laguna Madre, Corpus Christi Bay | 2011-2018 | Marine Science Institute, UT | Texas Seagrass Monitoring Program | Dataset | The University of Texas at Austin | NA | [Website](https://texasseagrass.org/) |
| 35 | USA | Texas | San Antonio | 2018 | Hobson, C., Whisenant, A. | Seagrass monitoring in San Antonio Bay, Texas with implications for management | Journal Article | Texas J. Sci. | 70: 1–14 | [10.32011/txjsci_70_1_Article1](https://doi.org/10.32011/txjsci_70_1_Article1) |
| 36 | USA | NA | US Gulf of Mexico | 2007 | Handley et al. | Seagrass status and trends in the Northern Gulf of Mexico: 1940 – 2002 | Book | US Geological Survey | NA | NA |

**Table S3.** Continued

| #N | Country | State | Location | Year | Author(s) | Title | Type | Publisher | Vol: pp | URL/doi |
| --- | --- | --- | --- | --- | --- | --- | --- | --- | --- | --- |
| 37 | USA | NA | US Gulf of Mexico | 2020 | Handley, L., Lockwood, C. | Seagrass status and trends update for the Northern Gulf of Mexico: 2002–2017 | Report | Gulf of Mexico Alliance | NA | NA |
| 38 | USA | NA | US Gulf of Mexico | 2021 | Darnell et al. | Seed reserve hot spots for the sub-tropical seagrass *Halodule wrightii* (shoal grass) in the Northern Gulf of Mexico | Journal Article | Estuaries and Coasts | 44: 339–351 | [10.1007/s12237-020-00808-y](https://doi.org/10.1007/s12237-020-00808-y) |
| 39 | USA; Mexico | Texas; Tamaulipas | Laguna Madre | 2002 | Tunnell, J.W., Judd, F.W. | The Laguna Madre of Texas and Tamaulipas | Book | Texas A&M University Press | NA | NA |
| 40 | Gulf of Mexico | NA | Gulf of Mexico | 2019 | Thorhaug et al. | Gulf of Mexico estuarine blue carbon stock, extent and flux: Mangroves, marshes, and seagrasses: A North American hotspot | Journal Article | Sci. Total Environ. | 653: 1253–1261 | [10.1016/j.scitotenv.2018.10.011](https://doi.org/10.1016/j.scitotenv.2018.10.011) |
| 41 | Mexico | Campeche | Laguna Términos | 2015 | Coria-Monter E, Durán-Campos E | Proximal analysis of seagrass species from Laguna de Términos, Mexico | Journal Article | Hidrobiologica | 25: 249–255 | NA |
| 42 | Mexico | Campeche | Laguna Términos | 2021 | Cuevas et al. | Spatial configuration of seagrass community attributes in a stressed coastal lagoon, southeastern Gulf of Mexico | Journal Article | Reg. Stud. Mar. Sci. | 48: 102049 | [10.1016/j.rsma.2021.102049](https://doi.org/10.1016/j.rsma.2021.102049) |
| 43 | Mexico | Campeche | Petenes | 2019 | Perez et al. | Distribución espacial de la vegetación acuática sumergida en los Petenes, Campeche | Journal Article | Terra Digitalis | 3(2): 1–11 | [10.22201/igg.25940694.2019.2.56](https://doi.org/10.22201/igg.25940694.2019.2.56) |
| 44 | Mexico | Campeche | Petenes | 2019 | Rosas-Valdez et al. | Seagrass and Fish in Los Petenes Biosphere Reserve, Campeche, Mexico: Spatial and Temporal Biomass Patterns | Journal Article | Thalassas | 35: 577–586 | [10.1007/s41208-019-00166-y](https://doi.org/10.1007/s41208-019-00166-y) |
| 45 | Mexico | Campeche | Petenes | 2021 | Cota TC, Herrera-Silveira JA | Seagrass contribution to blue carbon in a shallow karstic coastal area of the Gulf of Mexico | Journal Article | PeerJ | 9: e12109 | [10.7717/peerj.12109](https://doi.org/10.7717/peerj.12109) |
| 46 | Mexico | Campeche; Tamaulipas; Veracruz | Laguna Madre; Laguna Términos; Laguna Tamiahua; Laguna Alvarado | 2000 | Raz-Guzman, A., Barba, E. | Seagrass biomass, distribution and associated macrofauna in southwestern Gulf of Mexico coastal lagoons | Conference Proceedings | Proceedings Fourth International Seagrass Biology Workshop | 271–274 | NA |
| 47 | Mexico | Quintana Roo | Banco Chinchorro | 2009 | Cepeda-González et al. | Planeación para la conservación de la reserva de la biosfera Banco Chinchorro: Un esfuerzo conjunto | Report | The Nature Conservancy |  | [10.13140/2.1.4753.6642](https://doi.org/10.13140/2.1.4753.6642) |
| 48 | Mexico | Quintana Roo | Benito Juarez | 2021 | Hedley et al. | Seagrass depth distribution mirrors coastal development in the Mexican Caribbean – An automated analysis of 800 satellite images | Journal Article | Front. Mar. Sci. | 8: 1–24 | [10.3389/fmars.2021.733169](https://doi.org/10.3389/fmars.2021.733169) |

**Table S3.** Continued

| #N | Country | State | Location | Year | Author(s) | Title | Type | Publisher | Vol: pp | URL/doi |
| --- | --- | --- | --- | --- | --- | --- | --- | --- | --- | --- |
| 49 | Mexico | Quintana Roo | Puerto Morelos | 2013 | Zapata-Ramírez et al. | Accuracy of IKONOS for mapping benthic coral-reef habitats: A case study from the Puerto Morelos Reef National Park, Mexico | Journal Article | Int. J. Remote Sens. | 34: 3671–3687 | [10.1080/01431161.2012.716922](https://doi.org/10.1080/01431161.2012.716922) |
| 50 | Mexico | Tamaulipas | Laguna Madre | 2019 | Arellano-Méndez et al. | Structural complexity of tropical seagrasses meadows in a temperate lagoon in the Gulf of Mexico. A landscape ecology approach | Journal Article | J. Coast. Conserv. | 23: 969–976 | [10.1007/s11852-019-00701-2](https://doi.org/10.1007/s11852-019-00701-2) |
| 51 | Mexico | Tamaulipas | Laguna Madre | 2021 | DUMAC | Investigando el estado de los pastos marinos en la Laguna Madre | Website | DUMAC |  | [Website](https://dumac.org/en/2018/04/23/investigando-el-estado-de-los-pastos-marinos-en-la-laguna-madre/) |
| 52 | Mexico | Tamaulipas | Laguna Madre; Laguna Términos | 2005 | Barba et al. | Distribution patterns of estuarine caridean shrimps in the southwestern Gulf of Mexico | Journal Article | Crustaceana | 78: 709–726 | [10.1163/156854005774353502](https://doi.org/10.1163/156854005774353502) |
| 53 | Mexico | Tamaulipas | Laguna Morales | 2009 | García-Soriano et al. | Sitios de manglar con relevancia biológica y con necesidades de rehabilitación ecológica. Ficha de caracterización | Report | CONABIO |  | [Website](http://www.conabio.gob.mx/conocimiento/manglares/doctos/caracterizacion/GM44_Laguna_de_Morales_caracterizacion.pdf) |
| 54 | Mexico | Veracruz | Alvarado Lagoon | 2007 | Winfield et al. | ¿Controla la biomasa de pastos marinos la densidad de los peracaridos (Crustacea: Peracarida) en lagunas tropicales? | Journal Article | Rev. Biol. Trop. | 55: 43–53 | NA |
| 55 | Mexico | Veracruz | Alvarado Lagoon | 2014 | Rivera-Guzmán et al. | Long term state of coastal lagoons in Veracruz, Mexico: Effects of land use changes in watersheds on seagrasses habitats | Journal Article | Ocean Coast. Manag. | 87: 30–39 | [10.1016/j.ocecoaman.2013.10.007](https://doi.org/10.1016/j.ocecoaman.2013.10.007) |
| 56 | Mexico | Veracruz | Laguna Tamiahua | 1994 | Martínez-Murillo et al. | Ciliados asociados al pasto marino *Halodule beaudettei* en la laguna de Tamiahua, Veracruz, México | Journal Article | An. del Inst. Biol. la Univ. Nac. Autónoma México | 65: 11–18 | NA |
| 57 | Mexico | Veracruz | Laguna Tamiahua | 2000 | Díaz-Ruiz et al. | Distribution and abundance of *Syngnathus louisianae* and *Syngnathus scovelli* (Syngnathidae) in Tamiahua Lagoon, Gulf of Mexico | Journal Article | Ciencias Mar. | 26: 125–143 | [10.7773/cm.v26i1.567](https://doi.org/10.7773/cm.v26i1.567) |
| 58 | Mexico | Veracruz | Veracruz Reef System | 2008 | Terrados et al. | State of *Thalassia testudinum* Banks ex König meadows in the Veracruz Reef System, Veracruz, México | Journal Article | Aquat. Bot. | 88: 17–26 | [10.1016/j.aquabot.2007.08.003](https://doi.org/10.1016/j.aquabot.2007.08.003) |
| 59 | Mexico | Veracruz | Veracruz Reef System | 2011 | Terrados et al. | Cover and edge length to area ratio of seagrass (*Thalassia testudinum*) meadows in coral reef lagoons (Veracruz Reef System, Southwest Gulf of México) | Journal Article | Aquat. Conserv. Mar. Freshw. Ecosyst. | 21: 224–230 | [10.1002/aqc.1188](https://doi.org/10.1002/aqc.1188) |

**Table S3.** Continued

| #N | Country | State | Location | Year | Author(s) | Title | Type | Publisher | Vol: pp | URL/doi |
| --- | --- | --- | --- | --- | --- | --- | --- | --- | --- | --- |
| 60 | Mexico | Veracruz | Veracruz Reef System | 2016 | Arellano-Méndez et al. | Distribución espacial y estructura morfométrica de las praderas de *Thalassia testudinum* (Hydrocharitaceae) en dos arrecifes del Parque Nacional Sistema Arrecifal Veracruzano, México | Journal Article | Rev. Biol. Trop. | 64: 427–448 | [10.15517/rbt.v64i2.19810](https://doi.org/10.15517/rbt.v64i2.19810) |
| 61 | Mexico | Yucatan | Yucatán Coast | 2017 | Betzabeth PJE, de los Ángeles LCM | Spatial diversity of a coastal seascape: Characterization, analysis and application for conservation | Journal Article | Ocean. Coast. Manag. | 136: 185–195 | [10.1016/j.ocecoaman.2016.12.002](https://doi.org/10.1016/j.ocecoaman.2016.12.002) |
| 62 | Mexico | NA | Mexican Atlantic | 2018 | CONABIO | Benthonic maps | Dataset | CONABIO |  | [Website](http://www.conabio.gob.mx/informacion/gis/) |
| 63 | Mexico | NA | Mexican Caribbean | 2021 | Garcés-Cuartas et al. | Isotopic composition of aquatic and semiaquatic plants from the Mexican Caribbean: A baseline for regional ecological studies | Journal Article | Estuar. Coast. Shelf. Sci. | 260: 107489 | [10.1016/j.ecss.2021.107489](https://doi.org/10.1016/j.ecss.2021.107489) |
| 64 | Mexico | NA | National | 2018 | Herrera-Silveira et al. | Base de datos de almacenes de carbono en los pastos marinos de México. | Journal Article | Elem. Para Políticas Públicas | 2: 45–52 | NA |
| 65 | Mexico | NA | National | 2019 | Herrera-Silveira et al. | Pastos marinos, in: Estado del Ciclo del Carbono en México: Agenda Azul y Verde | Book Chapter | Programa Mexicano del Carbono | 150–177 | NA |
| 66 | Mexico | NA | Southern Gulf of Mexico | 2020 | Gallegos, M., Hernández, G. | Atlas de línea base ambiental del Golfo de México. Tomo VI: Pastos marinos | Book | CICESE |  | [Website](https://atlascigom.cicese.mx/es/dataset/libro-atlas-tomo-6) |
| 67 | Cuba | NA | Archipelago Sabana-Camagüey | 2007 | Martínez-Daranas B | Características y estado de conservación de los pastos marinos en áreas de interés del archipiélago Sabana-Camagüey, Cuba | Thesis | Universidad de La Habana |  | <https://aquadocs.org/items/1c80ffa0-9c3a-47ba-911b-3007cba5ef06> |
| 68 | Cuba | NA | Archipelago Sabana-Camagüey | 2021 | Martínez-Daranas et al. | Influence of several stressful factors on the condition of seagrasses at Sabana–Camagüey Archipelago, Cuba | Journal Article | Reg. Stud. Mar. Sci. | 47: 101939 | [10.1016/j.rsma.2021.101939](https://doi.org/10.1016/j.rsma.2021.101939) |
| 69 | Cuba | Sancti Spíritus | Bahía Buena Vista | 2021 | Martínez-Daranas et al. | Los pastos marinos del Parque Nacional Caguanes, Sancti Spíritus, Cuba | Journal Article | Rev. Investig. Mar. | 41: 28–46 | <https://revistas.uh.cu/rim/article/view/5280> |
| 70 | Cuba | Cienfuegos | Bahía Cienfuegos | 2009 | Moreira et al. | El impacto del huracán Dennis sobre el macrofitobentos de la Bahía de Cienfuegos, Cuba | Journal Article | Rev. Investig. Mar. | 30: 175–185 | NA |
| 71 | Cuba | Ciego de Ávila | Ciego de Ávila Norte | 2015 | Bustamante et al. | Caracterización de los pastos marinos de Paredón Grande, norte de la Provincia de Ciego de Ávila, Cuba | Journal Article | Rev. Investig. Mar. | 35: 74–90 | NA |
| 72 | Cuba | Ciego de Ávila | Ciego de Ávila Norte | 2019 | Cruz et al. | Estado de pastos marinos en dos playas de la cayería norte, antes y después del paso del huracán Irma por la provincia de Ciego de Ávila, Cuba | Journal Article | Rev. Ciencias Mar. y Costeras | 11: 85–108 | [10.15359/revmar.11-2.5](https://doi.org/10.15359/revmar.11-2.5) |

**Table S3.** Continued

| #N | Country | State | Location | Year | Author(s) | Title | Type | Publisher | Vol: pp | URL/doi |
| --- | --- | --- | --- | --- | --- | --- | --- | --- | --- | --- |
| 73 | Cuba | Ciego de Ávila | Ciego de Ávila Norte | 2020 | Cabrera et al. | Comparative growth and demographics of *Thalassia testudinum* meadows in Cuba using direct and reconstructive methods approaches to inform conservation efforts | Journal Article | Int. J. Recent Sci. Res | 11: 37446–37452 | [10.24327/IJRSR](https://doi.org/10.24327/IJRSR) |
| 74 | Cuba | Ciego de Ávila | Ana María Gulf | 2012 | Ventura Y, Rodríguez Y | Hábitats del golfo de Ana María identificados mediante el empleo de procesamiento digital de imágenes | Journal Article | Rev. Investig. Mar. | 32: 1–8 | NA |
| 75 | Cuba | Isla de la Juventud | Batabanó Gulf | 2008 | Cerdeira-Estrada et al. | Mapping of the spatial distribution of benthic habitats in the Gulf of Batabanó using Landsat-7 images | Journal Article | Ciencias Mar. | 34: 213–222 | [10.7773/cm.v34i2.1293](https://doi.org/10.7773/cm.v34i2.1293) |
| 76 | Cuba | Isla de la Juventud | Batabanó Gulf | 2008 | Arias-Schreiber et al. | Changes in benthic assemblages of the Gulf of Batabanó (Cuba) - Results from cruises undertaken during 1981-85 and 2003-04 | Journal Article | Panam. J. Aquat. Sci. | 3: 49–60 | <http://www.panamjas.org/pdf_artigos/PANAMJAS_3(1)_49-60.pdf> |
| 77 | Cuba | Pinar del Río | Guanahacabibes Gulf | 2013 | Cobián et al. | Caracterizacón de los ecosistemas costeros al norte del Área Protegida de Recursos Manejados Península de Guanahacabibes, Cuba | Journal Article | Rev. Ciencias Mar. y Costeras | 5: 37–55 | <https://doi.org/10.15359/revmar.5.3> |
| 78 | Cuba | Pinar del Río | Guanahacabibes Gulf | 2017 | Torres-Conde, E.G., Martínez-Daranas, B. | Los pastos marinos del Golfo de Guanahacabibes, Pinar del Río, Cuba | Journal Article | Rev. Investig. Mar. | 37: 1–15 | <https://revistas.uh.cu/rim/article/view/5001> |
| 79 | Cuba | Pinar del Río | Golfo Guanahacabibes | 2021 | Buesa, R.J. | Macrophytobenthos biomass in the Northwestern Cuban shelf, almost 50 years ago | Journal Article | Aquat. Bot. | 170: 103355 | [10.1016/j.aquabot.2021.103355](https://doi.org/10.1016/j.aquabot.2021.103355) |
| 80 | Cuba | Holguin | Guardalavaca | 2006 | Zayas et al. | Abundancia y diversidad de especies del fitobentos de playa Guardalavaca | Journal Article | Rev. Investig. Mar. | 27: 87–93 | NA |
| 81 | Cuba | Camagüey | Jardines de la Reina | 2018 | Bustamante et al. | Pastos marinos de Pasa Caballones, Parque Nacional Jardines de la Reina, Cuba | Journal Article | Rev. Investig. Mar. | 38: 28–44 | <https://revistas.uh.cu/rim/article/view/4182> |
| 82 | Cuba | NA | National | 2002 | Martínez-Daranas B | Variaciones morfológicas de *Halodule wrightii* Ascherson (Cymodoceaceae) en Cuba | Journal Article | Oceánides | 17: 93–101 | NA |
| 83 | Cuba | NA | National | 2011 | Guimarais et al. | Distribución de la familia Ruppiaceae en Cuba, nuevas consideraciones para su actualización en la flora del país | Journal Article | Mesoamericana | 15: 25–31 | NA |
| 84 | Cuba | NA | National | 2013 | Martínez-Daranas et al. | Distribución de *Halophila engelmanni* Ascherson (Hydrocharitaceae) en Cuba | Journal Article | Rev. Investig. Mar. | 33: 21–27 | NA |
| 85 | Cuba | NA | National | 2018 | Martínez-Daranas, B., Suárez, A.M. | An overview of Cuban seagrasses | Journal Article | Bull. Mar. Sci. | 94: 269–282 | [10.5343/bms.2017.1014](https://doi.org/10.5343/bms.2017.1014) |
| 86 | Cuba | NA | Northern Cuba; Batabanó Gulf | 2009 | Martínez-Daranas et al. | Los pastos marinos de Cuba: estado de conservación y manejo | Journal Article | Ser. Ocean. | 5: 24–44 | NA |

**Table S3.** Continued

| #N | Country | State | Location | Year | Author(s) | Title | Type | Publisher | Vol: pp | URL/doi |
| --- | --- | --- | --- | --- | --- | --- | --- | --- | --- | --- |
| 87 | Cuba | NA | Northern Cuba; Batabanó Gulf | 2009 | Hernández et al. | Evaluación de las posibles afectaciones del cambio climático a la biodiversidad marina y costera de Cuba | Report | Instituto de Oceanología |  | NA |
| 88 | Cuba | Camagüey | Nuevitas Bay | 2009 | Martínez-Daranas et al. | Spatial and seasonal variability of *Thalassia testudinum* in Nuevitas Bay, Cuba | Journal Article | Rev. Ciencias Mar. y Costeras | 1: 9–27 | [10.15359/revmar.1.1](https://doi.org/10.15359/revmar.1.1) |
| 89 | Cuba | La Habana | Rincón de Guanabo | 2017 | Aguilera, L | Cartografía de la istribución espacial del pasto marion en el PNP “Rincón de Guanabo”, La Habana | Thesis | Universidad de La Habana |  | NA |
| 90 | Cuba | La Habana | Santa Fe | 2016 | Gómez E, Martínez-Daranas B | Caracterización del macrofitobentos de la Laguna Grande, Santa Fe, La Habana, Cuba | Journal Article | Rev. Investig. Mar. | 36: 1–15 | <https://revistas.uh.cu/rim/article/view/5519> |
| 91 | Cuba | Camagüey | Santa Lucía | 2016 | Reyes, L. | Distribución y conservación de los pastos marinos en la playa Santa Lucía, Camagüey, Cuba | Thesis | Universidad de La Habana |  | <https://aquadocs.org/items/9c3c9e88-e08d-42d6-baeb-59de7d90974c> |
| 92 | Global | NA | NA | 2021 | UNEP-WCMC; Short | Global distribution of seagrasses | Dataset | UNEP-WCMC |  | [Website](http://wcmc.io/odv_WCMC_013_014) |
| 93 | Cuba, Mexico, USA | NA | NA | 1984-2021 | GBIF | Seagrass ocurrences | Dataset | Global Biodiversity Information Facility |  | [Website](https://www.gbif.org/) |

**Table S4.** Characterization of 17 seagrass time series sites (1987–2021) distributed in five subregions (NWF: Northwest Florida, WF: West Florida, SF: South Florida, TX: Texas, MXC: Mexican Caribbean). The values of population density (1985–2000), precipitation (1987–2021), sea surface temperature (SST) (2003–2021), chlorophyll-a (2003–2021), and particulate organic matter (POC) (2003–2021) are annual averages of the respective dataset availability periods. The hurricane impacts are the cumulative events at each site during 1987–2021. The color gradient is shown per variable across sites.

| **Subregion** | **Site** | **Seagrass extent trend** | **Popul. Density**  **(inhab. 100 m pixel)** | **Precip.**  **(mm yr^-1^)** | **SST (°C)** | **Chlor-a**  **(mg m^-3^)** | **POC**  **(mg m^-3^)** | **Hurricane Impacts** | **MPA cover (%)** |
| --- | --- | --- | --- | --- | --- | --- | --- | --- | --- |
| NWF | Big Bend A | Stable | 0.1 | 1,423 | 23.3 | 7.9 | 680 | 1 | 0.0 |
| NWF | Big Bend A | Stable | 0.1 | 1,343 | 23.9 | 12.2 | 869 | 0 | 99.1 |
| NWF | Choctawhatchee Bay | Increasing | 2.0 | 1,629 | 24.2 | 6.2 | 594 | 0 | 26.8 |
| NWF | St. Joseph Bay | Stable | 0.2 | 1,521 | 23.7 | 5.6 | 555 | 2 | 78.4 |
| WF | Charlotte Harbor (North) | Stable | 1.8 | 1,356 | 26.2 | 4.3 | 433 | 1 | 99.5 |
| WF | Clearwater Harbor | Increasing | 11.9 | 1,249 | 25.1 | 3.7 | 435 | 0 | 99.7 |
| WF | Sarasota Bay | Stable | 4.3 | 1,359 | 25.3 | 3.7 | 410 | 0 | 0.6 |
| WF | Springs Coast C | Stable | 2.4 | 1,354 | 24.3 | 3.8 | 432 | 0 | 4.3 |
| WF | St. Joseph Sound | Increasing | 8.9 | 1,333 | 24.7 | 4.1 | 456 | 0 | 99.9 |
| WF | Tampa Bay | Increasing | 7.5 | 1,337 | 25.3 | 6.3 | 555 | 0 | 73.2 |
| SF | Biscayne Bay | Decreasing | 15.3 | 1,437 | 27.0 | 0.5 | 110 | 2 | 95.9 |
| SF | Florida Bay (East) | Increasing | 0.2 | 1,356 | 27.2 | 3.9 | 367 | 2 | 98.4 |
| TX | Corpus Christi Bay | Stable | 2.7 | 748 | 26.2 | 3.6 | 398 | 0 | 1.0 |
| TX | Lower Laguna Madre | Stable | 0.1 | 631 | 27.4 | 2.4 | 322 | 3 | 8.0 |
| TX | Upper Laguna Madre | Stable | 0.4 | 687 | 27.7 | 3.1 | 387 | 2 | 30.5 |
| MXC | Benito Juarez | Increasing | 3.0 | 1,346 | 27.9 | 0.2 | 56 | 1 | 100.0 |
| MXC | Puerto Morelos | Increasing | 0.2 | 1,346 | 28.1 | 0.1 | 43 | 2 | 91.8 |
